# Evolutionary regain of lost gene circuit function

**DOI:** 10.1101/804187

**Authors:** Mirna Kheir Gouda, Michael Manhart, Gábor Balázsi

**Author notes:** To whom correspondence should be addressed: Gábor Balázsi, Ph.D., The Louis and Beatrice Laufer Center for Physical & Quantitative Biology, 115C Laufer Center, Z-5252, Stony Brook University, Stony Brook, NY 11794.

## Abstract

Evolutionary reversibility - the ability to regain a lost function - is an important problem both in evolutionary and synthetic biology, where repairing natural or synthetic systems broken by evolutionary processes may be valuable. Here, we use a synthetic positive-feedback (PF) gene circuit integrated into haploid *Saccharomyces cerevisiae* cells to test if the population can restore lost PF function. In previous evolution experiments, mutations in a gene eliminated the fitness costs of PF activation. Since PF activation also provides drug resistance, exposing such compromised or broken mutants to both drug and inducer should create selection pressure to regain drug resistance and possibly PF function. Indeed, evolving seven PF mutant strains in the presence of drug revealed three adaptation scenarios through genomic mutations outside of the PF circuit that elevate PF basal expression, possibly by affecting transcription, translation, degradation and other fundamental cell functions. Nonfunctional mutants gained drug resistance without ever developing high expression, while quasi-functional and dysfunctional PF mutants developed high expression which then diminished, although more slowly for dysfunctional mutants where revertant clones arose. These results highlight how intracellular context, such as the growth rate, can affect regulatory network dynamics and evolutionary dynamics, which has important consequences for understanding the evolution of drug resistance and developing future synthetic biology applications.

**Significance Statement:** Natural or synthetic genetic modules can lose their function over long-term evolution if the function is costly. How populations can evolve to restore broken functions is poorly understood. To test the reversibility of evolutionary breakdown, we use yeast cell populations with a chromosomally integrated synthetic gene circuit. In previous evolution experiments the gene circuit lost its costly function through various mutations. By exposing such mutant populations to conditions where regaining gene circuit function would be beneficial we find adaptation scenarios with or without repairing lost gene circuit function. These results are important for drug resistance or future synthetic biology applications where loss and regain of function play a significant role.

## 1. Introduction

Two ways for cells to survive stress and buy time until beneficial genetic alterations arise are through sensing and responding or through bet-hedging [1, 2]. Gene regulatory networks evolve to provide cells with sufficient stress-protective gene expression according to these strategies. While stress-protective mutations improve the chance of survival, a tradeoff often exists between the cost and benefit of such protective mechanisms [3]. For example, the expression of stress-protective genes can have a net cost in the absence of stress or even in stress if expression surpasses the levels necessary for survival [4–7]. Consequently, protective but costly gene function tends to diminish or vanish from the population in the absence of stress [5, 8–10]. How it might reappear again (evolutionary “reversal”) when the stress resumes [11, 12] is poorly understood, specifically for gene regulatory networks that lost their costly activity. Indeed, loss-of-function mutations occur widely in laboratory evolution experiments [5, 8, 13–16], suggesting this is a common mode of adaptation to a new environment. However, few experiments have tested how such lost functions could be restored.

Besides experimental studies of natural gene network evolution under controlled conditions [5, 8, 17–21], synthetic gene circuits can serve as well-characterized models of natural stress-response modules in evolution experiments [9, 22, 23]. Well-controlled and tunable synthetic gene circuits that interact minimally with the host genome [24–27] can aid the interpretation of experimental outcomes. Considering future applications of synthetic biology [28, 29], it is important to explore potential evolutionary strategies that can restore synthetic gene circuit function if it happens to break over time [9, 30, 31], without its reintroduction or repair by rational means or mutagenesis [32]. Directed evolution studies have improved enzymes or metabolic pathways [33, 34], but only through mutagenesis in single proteins, leaving it unclear whether noncoding or genomic mutations could improve or restore gene network performance. Adaptation under function-restoring selection pressure allowing host genome changes could reveal essential parameters and new methods to design and improve engineered cell fitness and robustness in various growth conditions.

To understand how a network that lost its costly activity in the absence of stress adapts and possibly regains function in the presence of stress, here we used previously evolved, broken versions of a synthetic “positive feedback” (PF) gene circuit originally integrated into the haploid *S. cerevisiae* YPH500 genome [6]. Many different rtTA mutants arose in previous evolution experiments [9] apparently eliminating costly rtTA function in the absence of antibiotic stress. We evolved seven such broken PF mutants in both inducer and antibiotic, where regaining rtTA function should be beneficial. By examining the phenotypic and genetic changes through fluorescence, fitness measurements, and sequencing, we observed three different classes of evolutionary dynamics, depending on whether ancestral mutants were quasi-functional, dysfunctional, or nonfunctional. In quasi-functional mutants, slow growth from drug exposure initially enriched the high expressor subpopulation, but then new drug resistance mutations slightly elevated basal expression, eliminating the benefit of high expression and diminishing the high-expressor fraction through a growth-dependent shift in dynamics. Nonfunctional mutants acquired new drug resistance mutations that slightly elevated basal expression and never developed high expression. Finally, the dysfunctional mutant populations evolved similarly to quasi-functional mutants, but more slowly and gave rise to clones with repaired network function. Overall, we found numerous extra-circuit mutations, but no novel coding sequence mutations inside the gene circuit directly related to these expression changes. Our findings provide insights into the evolutionary reactivation of broken network modules, depending on their dynamics as well as the costs and benefits after the stress recurs.

## 2. Results

### 2.1. Hyperinduction and slow growth reveal three classes of rtTA mutations

The original PF gene circuit [6] consists of a Doxycycline-inducible rtTA transcriptional activator that identically upregulates both its own expression and the expression of the Zeocin resistance gene *zeoR* fused to yEGFP (*yEGFP::zeoR*) by binding to two *tetO2* operator sites upstream of the *rtTA* and the *yEGFP::zeoR* coding regions (Figure 1A). The yEGFP::ZeoR bifunctional fusion protein [35] directly reports ZeoR levels and protects cells by directly binding to Zeocin [36] to prevent its intercalation into DNA, which causes DNA breaks and thus cell cycle arrest or death [37]. Previously, we evolved yeast populations carrying the PF circuit [9] in 2 μg/mL of Doxycycline inducer without Zeocin antibiotic, a condition denoted “D2Z0” where the first number represents Doxycycline concentration in μg/mL, while the second number represents Zeocin concentration in mg/mL. Since the activation of the PF stress response module is costly in the absence of stress, the rtTA protein apparently lost its function during these evolution experiments through different coding sequence mutations [9]. To test whether evolutionary repair of lost rtTA function was possible, we further evolved seven of these loss-of-function rtTA mutants from the previous experiment (Figure 1B) in the condition D2Z2 (2 µg/mL Doxycycline and 2 mg/mL Zeocin), which should activate the original PF gene circuit or any reverting mutants, ensuring their survival in stress. We will refer to the seven ancestral mutations by their genotype: “Missense 1, 2, 3, and 4” correspond to rtTA_+189C→G_, rtTA_+562T→C_, rtTA_+275G→A_, and rtTA_+13G→T_ respectively; “Nonsense” corresponds to rtTA_+442G→T_; “Duplication” corresponds to rtTA_+95, 30bp_; and “Deletion” corresponds to rtTA_+651, 78bpΔ_ (Table S1).

**Figure 1.**
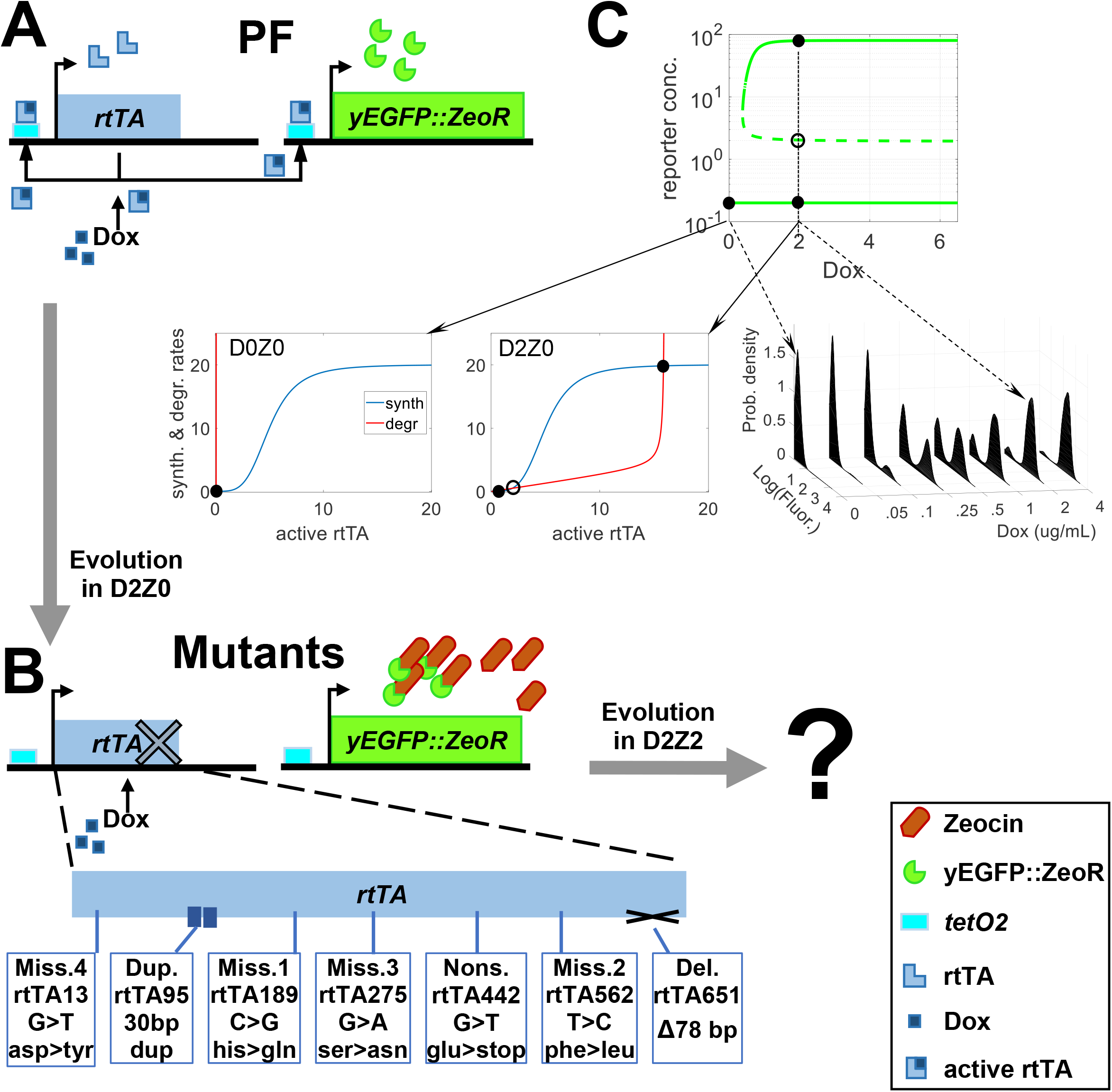
The PF gene circuit lost bistability and costly rtTA function by multiple mutations. **(A)** In the original PF gene circuit, the inducer Doxycycline (Dox) binds and activates the rtTA protein, which activates its own gene and *yEGFP::zeoR*. Since rtTA activity is costly, loss of rtTA function is evolutionarily beneficial in D2Z0. **(B)** We selected seven mutants that arose in D2Z0, improving fitness by PF breakdown. Now we evolve each mutant in D2Z2 where regaining rtTA function would be beneficial, since rtTA can activate expression of the yEGFP::ZeoR protein, which binds to Zeocin and prevents it from harming the cell. DxZy denotes the concentrations of the inducer **<u>D</u>**ox and the antibiotic **<u>Z</u>**eocin in μg/ml and mg/ml respectively. **(C)** The original PF gene circuit undergoes a saddle-node bifurcation, changing from monostable to bistable dynamics when Dox exceeds a threshold. The top graph shows yEGFP::zeoR levels versus Dox from a mathematical model (SI model). The blue and red curves on the bottom graphs represent rtTA synthesis and degradation rates; while filled and open circles denote stable and unstable steady states, respectively. The active rtTA levels corresponding to the circles impose the yEGFP::zeoR levels on the top graph and on the right, where experimental yEGFP::zeoR expression histograms versus Dox demonstrate the bifurcation.

To understand the dynamics of the original PF gene circuit (Figure 1A) and its seven rtTA mutants (Figure 1B), we studied a simple mathematical model of this genetic autoregulatory module (SI model). In rate-balance plots from these models (Figure 1C), intersections between rtTA synthesis (blue sigmoid) and loss (red elbow) rate curves correspond to stable (full circles) and unstable (open circles) cellular *rtTA* gene expression states that impose corresponding *yEGFP::zeoR* levels. For example, the original PF gene circuit [6] is monostable below a threshold concentration of ∼0.05 μg/mL Doxycycline, with a single intersection corresponding to a single low rtTA expression state imposing low *yEGFP::zeoR* expression. As Doxycycline concentrations increase, the elbow-shaped curve of unbound rtTA loss (from combined dilution, degradation, inducer binding) moves rightward, approaching the sigmoidal synthesis curve until they meet and intersect three times. Thus, for Doxycycline concentrations above the threshold, a second stable high expression state emerges, indicating a transition from monostable to bistable dynamics through a saddle-node bifurcation (Figure 1C). Losing the high gene expression peak during long-term evolution in D2Z0 [9] could indicate either that a mutant gene circuit became completely non-inducible, or that the sigmoidal rtTA synthesis rate curve shifted somehow, elevating the bistability threshold beyond D2Z0. Still, the elbow-shaped loss curve also slides rightward as Doxycycline increases, so it could still reach and intersect a right-shifted sigmoid three times, causing a high expression peak to still emerge at sufficiently high Doxycycline concentrations. To examine the possibility of such weaker but still present rtTA function, we looked for high expressors in flow cytometry histograms of each mutant in D2Z0. Indeed, upon closer examination, we noticed ∼1% high-expressing cells in Missense 1 and 2 populations, but not any other mutants (Figures S3-S8). Therefore, we deem the Missense 1 and 2 mutant gene circuits to be quasi-functional rather than completely nonfunctional (Table S1).

To fully test which of the seven loss-of-function PF mutants are still quasi-functional, we next hyper-induced them with excess Doxycycline. We expected that hyperinduction would shift the elbow-shaped curve of rtTA loss rightward and move quasi-functional rtTA mutants into their bistable regime (Figure 1C, SI model) thus generating high-expressor subpopulations, while nonfunctional mutants would remain unimodal. To this end we grew clonal populations carrying each rtTA mutation in D6Z0 (Figure 2A,B), a threefold higher (6 µg/mL) Doxycycline concentration than D2Z0 where they previously arose. These hyper-inducing conditions confirmed that Missense 1 and 2 were indeed quasi-functional, while the remaining five mutations - Missense 3 and 4, Nonsense, Duplication, and Deletion - still appeared uninducible and thus nonfunctional (Table S1). Excess Doxycycline did not alter the growth rate of any clones, indicating that hyperinduction is not toxic for any mutant. Thus, any further mutations abolishing leftover rtTA function in Missense 1 and 2 would not be beneficial. The results did not change upon hyper-inducing the mutants in 8 µg/mL Doxycycline (Figure S1A). Hysteresis experiments further confirmed the upshift of bistability range for Missense 1 and 2 compared to PF (Figure S12).

**Figure 2.**
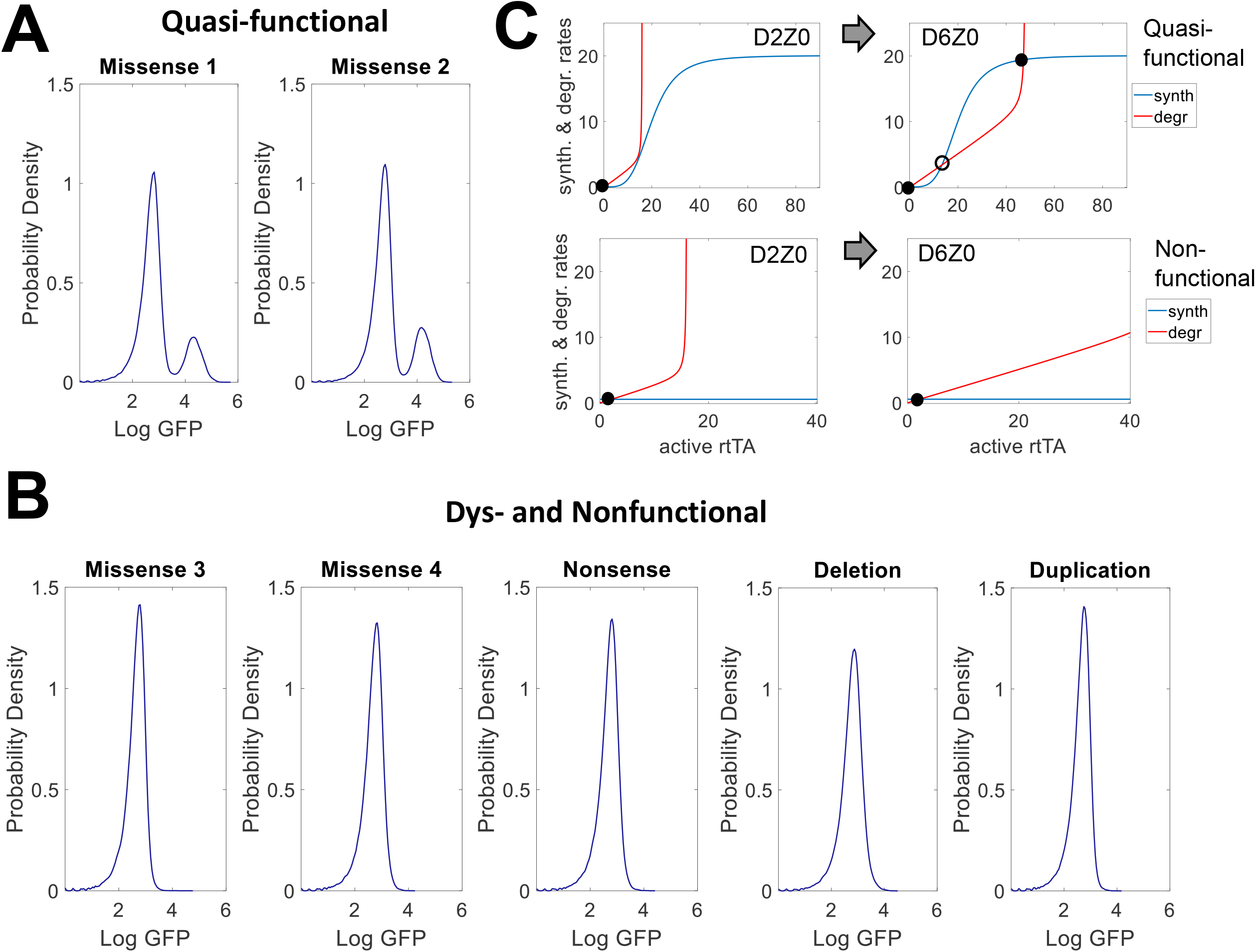
Quasi-functional rtTA mutants revealed by hyperinduction. **(A)** Gene expression histograms of quasi-functional PF mutants hyperinduced in D6Z0 (24h). A high expression peak in this condition indicates bistabiltiy. **(B)** Gene expression histograms of dysfunctional and nonfunctional PF mutants in D6Z0 (24h). The lack of a high expression peak in this condition indicates that they remain monostable even despite hyperinduction. **(C)** Plots illustrating the dynamical effect of hyperinduction on quasi-functional and nonfunctional PF mutants. Filled and open circles denote stable and unstable steady states, respectively.

Mathematical models suggested another mechanism besides hyperinduction that can cause stable high expression. Slow growth reduces dilution of cell contents and thus tilts downward the rtTA loss curve, besides shifting it rightward. This moves the quasi-functional PF mutants into the bistable regime (SI model), similar to growth-mediated bistability observed in other systems [38, 39]. To test this, we grew PF mutants as well as standard PF cells in D2Z0 with 7.5% ethanol that slowed the growth rate to a value similar to that in D2Z2 (Figures S3-S8). Interestingly, ethanol strongly enriched the high-expressor fraction of ancestral PF cells, from 71% to 91% in D2Z0. As expected, slow growth due to ethanol also enriched the high expressor fractions of Missense 1 (more than twofold) and Missense 2 (about ninefold). Most surprisingly, ethanol caused a few high expressor cells to appear even in Missense 3, which failed to respond to hyperinduction in D6Z0 or D8Z0. Therefore, we classify Missense 3 as a dysfunctional mutant (Table S1), unlike the nonfunctional mutants Nonsense and Deletion, which do not become bistable even in slow growth. We obtained comparable results using Cisplatin, which interferes with DNA [40, 41] like Zeocin. In contrast, G418 did not have this effect, presumably because protein synthesis inhibition [42] abolishes gene circuit function (Figures S3-S8). Overall, this data and the model indicate that slow growth due to stressors can cause shifts in dynamics that generate high expressors in Missense 3 and can enrich the already present high expressor subpopulations of Missense 1 and 2.

### 2.2. Evolution does not revert quasi-functional mutants

Next, we set out to test if evolution could revert the two quasi-functional mutants (Missense 1 and 2) to regain stronger rtTA function. Thus, we evolved three replicates each of Missense 1 and 2 in D2Z2 medium, where regaining high rtTA activity would be beneficial by shifting cells into the drug-resistant state of high yEGFP::ZeoR expression. For control purposes, we also propagated the same initial populations in D2Z0.

We observed qualitatively-similar evolutionary dynamics for both Missense 1 and Missense 2 (Figure 3), which started with a minimal (0.47% and 0.63%) subpopulation of high-expressor cells immediately upon transfer from D2Z0 into D2Z2. Early on, population fitness levels dropped significantly in all D2Z2 cultures compared to the control D2Z0 cultures (Figure 3B,D). Soon afterwards, drug exposure generated a substantial high-expressing subpopulation for a few days (Figure 3A,C). Growth rates started recovering in ∼ 4 days, as the high-expressor peak increased (Figure 3B,D). From day 1 onward, we also observed a slight upward fluorescence shift for low expressors, corresponding to elevated basal yEGFP::ZeoR levels compared to D2Z0 controls (Figure 3A,C). This slight upward expression shift is clearly distinguishable from “induced” high expression and resembles phenotypes arisen previously during evolution in Zeocin drug (D0Z2) [9], where various extra-circuit mutations, often coupled with intra-circuit synonymous and promoter mutations, elevated the low, basal yEGFP::ZeoR expression. Concurrently, the high-expressor population fraction reached a maximum around day 4 and then returned to the same low level as in D2Z0 within ∼ 8 days (Figure 3A,C).

**Figure 3.**
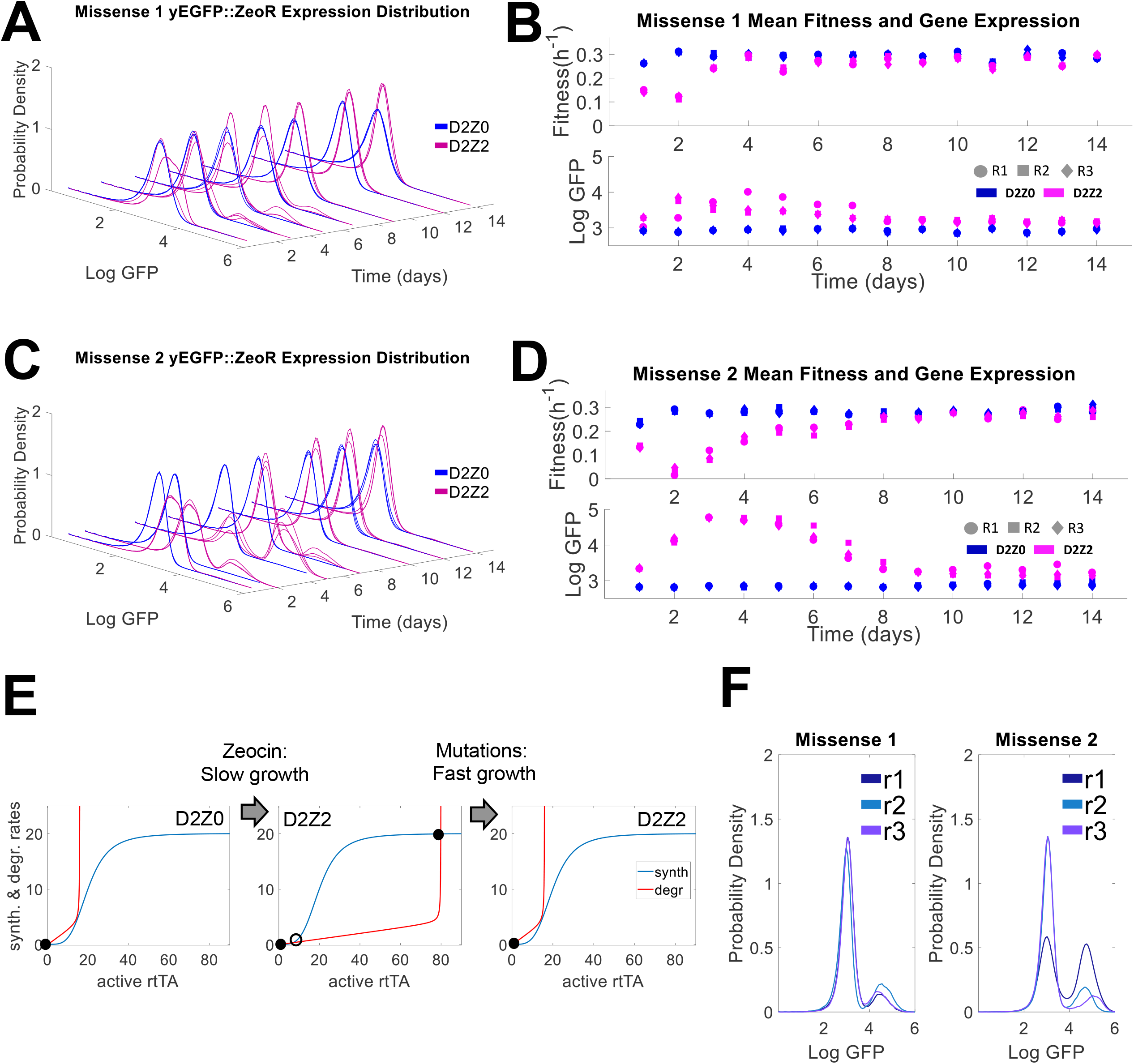
Evolutionary dynamics of quasi-functional mutants. **(A)** Histogram of *yEGFP::zeoR* expression for Missense 1 replicates in D2Z0 (blue, control) and D2Z2 (magenta) over the course of 14 days. High-expressor subpopulations emerge within the first two days (∼ 12 generations), then reach a peak and diminish within the next five days (∼ 30 generations). The low-expressor subpopulation shifts to slightly higher expression in D2Z2 compared to the mean expression in D2Z0. **(B)** Top: Population fitness (exponential growth rate) of Missense 1 cells in D2Z0 (blue) and D2Z2 (magenta) over the course of 14 days. Initially, fitness drops in D2Z2 cultures compared to D2Z0 cultures, but then the cells recover within ∼3 days and reach a fitness comparable to the D2Z0 cultures. Bottom: average gene expression levels corresponding to the histograms for Missense 1 in D2Z0 (blue) and D2Z2 (magenta). **(C)** Histogram of *yEGFP::zeoR* expression for Missense 2 replicates in D2Z0 (blue) and D2Z2 (magenta) over the course of 14 days. High-expressor subpopulations emerge within the first two days (∼ 12 generations), then reach a peak and diminish within the next six days (∼ 34 generations). Similar to missense mutant 1 cultures, the missense 2 dominant low-expressors population maintains a slightly higher GFP mean in D2Z2 compared to the mean expression in D2Z0. **(D)** Top: Population fitness (exponential growth rate) of Missense 2 in D2Z0 (blue) and D2Z2 (magenta) over the course of 14 days. Initially, fitness drops for D2Z2 cultures compared to D2Z0 cultures. Cells cultured in D2Z2 recover within a week and reach a fitness comparable to the D2Z0 cultures. Bottom: Mean *yEGFP::zeoR* expression of Missense 2 in D2Z0 (blue) and D2Z2 (magenta) corresponding to the histograms for Missense 2 in D2Z0 (blue) and D2Z2 (magenta). **(E)** Changes of PF dynamics due to slower growth and selection in Zeocin, and then, reversion to normal growth in D2Z2 after the emergence of drug resistance mutations. Filled and open circles denote stable and unstable steady states, respectively. **(F)** Gene expression histograms of evolved Missense 1 and Missense 2 population replicates hyperinduced in D6Z0 after the end of experimental evolution. A high expression peak in this condition characterizes the presence of quasi-functional mutants in the population.

Considering that high expressors were rare in D2Z0, while in D2Z2 their fraction initially increased and then later dropped back to the D2Z0 level, at least two hypotheses are possible. First, compensatory mutations in rtTA or elsewhere could improve rtTA activator function, thereby increasing the high expressor fraction, and then later subsequent mutations could revert this effect. To identify such compensatory mutations, we performed whole-genome sequencing (WGS) on one replicate population each for both mutants on days 1, 3, and 14. The original rtTA mutations already present at the beginning of the experiment were identified at 100% frequency at all time points for both Missense 1 and Missense 2. However, we identified no other intra-circuit mutations. Since certain variant types (e.g., deletions and duplications) in the PF gene circuit are difficult to detect by WGS [24], we also performed Sanger sequencing of the PF region in 10 individual clones from each mutant at day 14 (Table S2). All Missense 1 clones carried a deletion of the first *tetO2* operator upstream of the *yEGFP::zeoR* coding region, whereas the other *tetO2* site stayed intact. All Missense 2 clones carried only their original gene circuit mutation without any *tetO2* or other circuit modifications, despite their identical phenotypes to Missense 1. Therefore, there is no phenotypic signature directly and specifically attributable to the *tetO2* operator site deletion (Table S3). Overall, the only *de novo* intra-circuit mutation we found did not explain the observed gene expression changes.

High expression requires rtTA function. How then could a substantial high-expressor peak emerge and later diminish without any changes in the rtTA coding sequence or its promoter? Modeling and experiments suggested a second hypothesis: by decelerating growth and reducing dilution, Zeocin tilts downward and shifts rightward the red elbow curve of rtTA loss (Figure 3E). This causes a shift in dynamics that enriches the rare high-expressing drug resistant cells that preexist in D2Z0 (SI model), which are then further enriched by phenotypic selection. At the same time, the low-expressing subpopulation will remain drug-sensitive, enabling any mutants that elevate basal expression to grow faster and spread in the population. Once such fast-growing mutants arise, their faster growth/dilution rate tilts upward and shifts leftward the red elbow curve (Figure 3E), causing a return of dynamics towards monostability (SI model). Concurrently, the benefits of high-expression also vanish, causing the high-expressor fraction to diminish back to the D2Z0 levels. Importantly, these events can occur without any mutations in the PF gene circuit, as long as extra-circuit mutations can raise basal yEGFP::ZeoR expression to promote fast growth in Zeocin.

To identify such extra-circuit mechanisms of *yEGFP::zeoR*-dependent Zeocin resistance, we looked for genomic, extra-circuit mutations in the WGS data (Materials and Methods). In the Missense 1 population at day 14 we found moderate-frequency (∼14-15%), nonsynonymous mutations in two genes affecting protein stability and transcription, PAH1 and SET3; including a 1 bp frameshift deletion in the latter. Additional mutations in other genes and intergenic regions exist at lower frequencies (Dataset S1). In contrast with Missense 1, evolved Missense 2 populations carried multiple extra-circuit mutations exceeding 90% in frequency at day 14 (Dataset S2). Mutations in the genes PHB2, MDM32, and COX1 suggest alteration in mitochondrial metabolism and function. We also observed a synonymous mutation in the FG-nucleoporin NUP159 gene, which is involved in post-transcriptional regulation [43].

To test if elevated *yEGFP::zeoR* gene expression mediates drug resistance without rtTA activity, we compared the growth of Missense 1 and 2 evolved populations and clonal isolates to their unevolved Missense 1 and 2 ancestors in Zeocin only media (D0Z2). Evolved Missense 1 and 2 populations and clonal isolates grew significantly faster than the ancestral PF or unevolved Missense 1 and 2 cells in D0Z2 (Figure S9), indicating that the evolved Missense 1 and 2 populations were Zeocin-resistant independently of rtTA.

If the rtTA coding sequence and PF gene circuit dynamics did not change during evolution, hyperinduction should affect Missense 1 and 2 similarly to their ancestors. To test this, we hyper-induced each final evolved Missense 1 and 2 replicate population in D6Z0. Indeed, the evolved Missense 1 and 2 populations responded to D6Z0 (Figure 3F) and to D8Z0 (Figure S1) as their ancestors did. Likewise, hyperinduction did not affect the growth rates of the evolved populations. Overall, the Missense 1 and 2 evolution experiments indicate that slow growth and phenotypic selection initially enrich the high-expressor fraction, but then extra-circuit mutations accelerate growth by elevating basal stress-protective yEGFP::ZeoR expression, which shifts the dynamics to diminish the high-expressor fractions back to levels equivalent to D2Z0.

### 2.3. Most nonfunctional mutants never regain rtTA function

As the lack of high expression in D6Z0 (Figure 2) indicates, the four other initial PF mutants (Missense 4, Nonsense, Duplication, and Deletion) had mutations that disrupted rtTA protein function such that it became completely uninducible, regardless of inducer amount (Figure 2) or growth rate (Figures S3-S8). To test if evolution could restore rtTA function in any of these nonfunctional mutants, we also evolved three replicates of each in D2Z2 where regaining rtTA activity would generate a beneficial high *yEGFP::ZeoR* expression peak.

Early in the evolution experiment (Figure 4A-H), the growth rate in D2Z2 of each nonfunctional mutant population dropped significantly below the controls evolving in D2Z0. However, the growth rates of the Missense 4, Nonsense and Duplication populations then recovered over ∼ 5 days, while the growth rates of the Deletion populations recovered within 3 days, approaching that of D2Z0 control cultures. Unlike quasi-functional mutants, nonfunctional mutants never gave rise to a high-expressor subpopulation (Figure 4A-D). The *yEGFP::zeoR* expression distributions remained unimodal but shifted slightly upwards compared to the basal (D2Z0 control) expression in all experimental cultures. Consequently, these evolving cell populations relied again on *yEGFP::zeoR* to gain drug resistance and increase their fitness, without repairing rtTA function. Indeed, Sanger sequencing of 10 isolated clones from each mutant revealed no additional mutations inside the gene circuit, except for the deletion of the first *tetO2* site upstream of *yEGFP::zeoR* in 0/10 Missense 4 clones, 1/10 Nonsense clones, 8/10 Deletion clones, and 2/10 Duplication clones (Table S4). All mutants retained the original rtTA mutation and no other rtTA mutations were detected. As with Missense 1 and 2, there was no phenotype associated with the *tetO2* deletion (Table S4).

**Figure 4.**
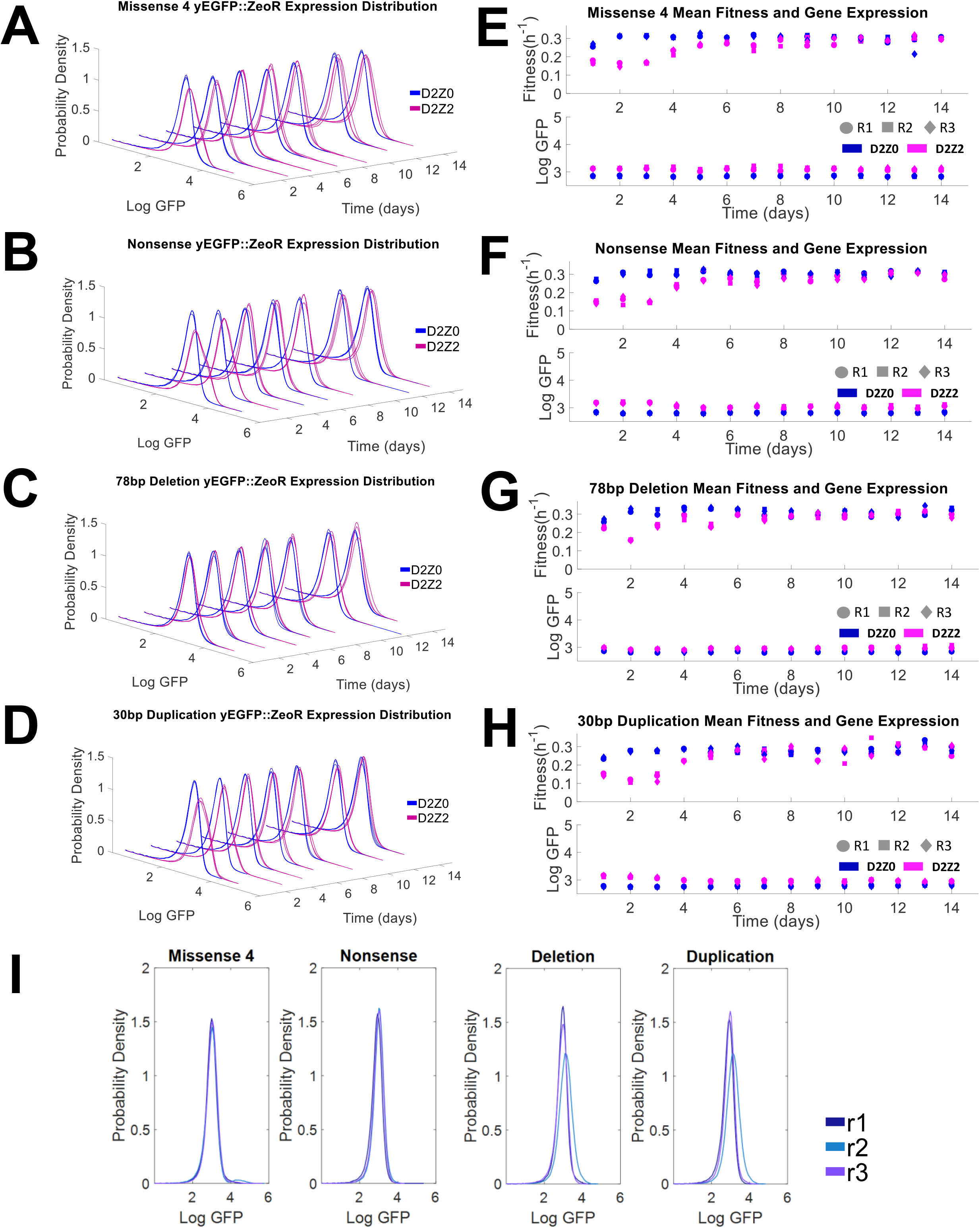
Evolutionary dynamics of nonfunctional mutants that never regain rtTA function. **(A-D)** Histograms of *yEGFP::zeoR* expression of 3 replicates of Missense 4, Nonsense, Deletion and Duplication mutants, respectively, in D2Z0 (blue) and D2Z2 (magenta) over the course of 14 days. These mutants do never exhibit any bimodality while evolving in D2Z2, but they develop a slightly higher basal yEGFP::zeoR mean in D2Z2 compared to the mean expression in D2Z0. **(E-H)** Fitness (Left) and mean *yEGFP::zeoR* expression (Right) plots for Missense 4, Nonsense, Deletion and Duplication mutants, respectively, computed for each day during the experiment. Early in the experiment, we observe a drop in fitness of D2Z2 cultures compared to D2Z0 cultures. Cells cultured in D2Z2 recover within ∼4 days and reach a fitness level comparable to the D2Z0 cultures. Additionally, D2Z2 cultures maintain a higher GFP expression mean compared to D2Z0 cultures for all presented mutants. **(I)** Gene expression histograms of evolved population replicates hyperinduced in D6Z0 after the end of experimental evolution.

To identify possible extra-circuit mutations causing the slight basal expression increase, we analyzed the WGS data. We found mutations in or near genes controlling mRNA and protein levels through degradation or synthesis, such as the poly-A tail shortening CCR4, the TFIID subunit TAF2, the ribosomal subunit RPL41A, and others (Datasets S4-S7).

To examine rtTA nonfunctionality at the end of the evolution experiment, we hyper-induced final evolved populations in D6Z0 (Figure 4I). As expected, the yEGFP::ZeoR expression of nonfunctional mutants remained unimodal and low in D6Z0 (Figure 4I) and in D8Z0 (Figure S1), indicating that rtTA and the gene circuit in these mutants remained nonfunctional at the end of evolution in D2Z2, as anticipated.

As we did for Missense 1 and 2, we also examined rtTA-independent Zeocin resistance of nonfunctional mutants by growing evolved populations and isolated clones in D0Z2 media for 24 hours. The evolved Missense 4, Nonsense, and Duplication populations and the isolated clones grew significantly faster than their unevolved ancestors (Figure S9). Interestingly, the unevolved Deletion mutant seemed to have some preexisting yEGFP::ZeoR-dependent resistance to Zeocin (Figure S9). This corroborates the faster Deletion recovery and higher yEGFP::ZeoR basal expression compared to all other non-functional mutants upon Zeocin exposure, already at day 1 (Figure S11), and reinforces general reliance on slightly elevated yEGFP::ZeoR expression for drug resistance.

### 2.4. Dysfunctional mutants can regain PF gene circuit function

Surprisingly, during evolution in D2Z2, all three replicates of the dysfunctional rtTA mutant (Missense 3) temporarily gave rise to a significant high-expressor subpopulation that persisted for over a week for each replicate. Ultimately, towards the end of the evolution experiment, these high-expressor subpopulations diminished and the fluorescence shift of the low peak indicated a slight increase in basal yEGFP::zeoR expression similar to all other genotypes. The fitness trends of each Missense 3 replicate resembled those of the other mutants: the initially low population fitness levels recovered to normal (control) levels within ∼ 4 days (Figure 5A,B).

**Figure 5.**
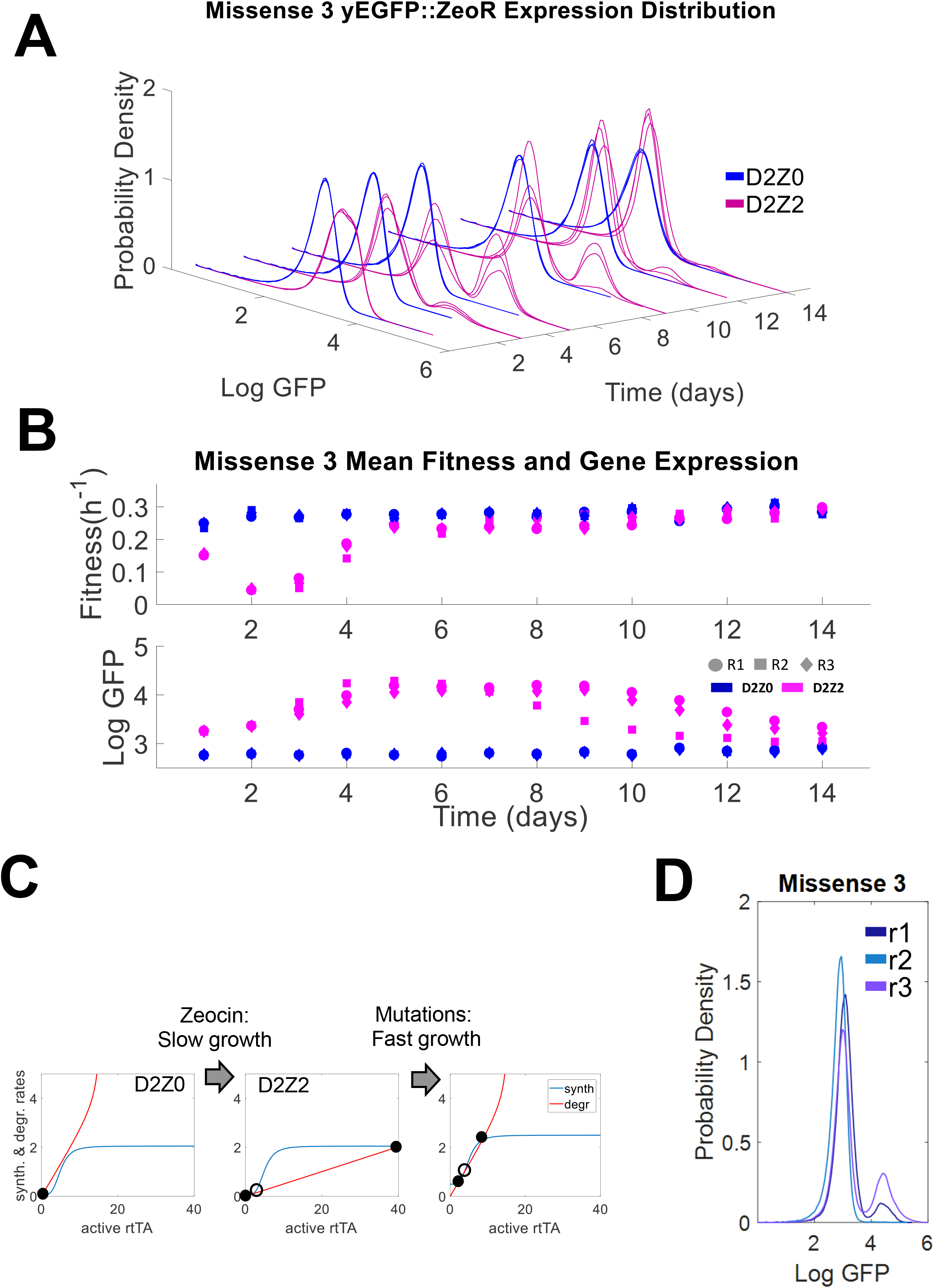
Evolutionary dynamics of the dysfunctional mutant Missense 3. **(A)** Histogram of *yEGFP::zeoR* expression for Missense 3 replicates in D2Z0 (blue) and D2Z2 (magenta) over the course of 14 days. High-expressor subpopulations emerge within the first three days (∼ 18 generations), then reach a peak and diminish within the next 11 days (∼ 80 generations). The low-expressor subpopulation shifts to slightly higher expression in D2Z2 compared to the mean expression in D2Z0. **(B)** Top: Population fitness (exponential growth rate) of Missense 3 cells in D2Z0 (blue) and D2Z2 (magenta) over the course of 14 days. Initially, fitness drops in D2Z2 cultures compared to D2Z0 cultures, but then the cells recover within ∼3 days and reach a fitness comparable to the D2Z0 cultures. Bottom: average gene expression levels corresponding to the histograms for Missense 1 in D2Z0 (blue) and D2Z2 (magenta). **(C)** Evolutionary changes in gene circuit dynamics involve slow growth-induced bistability due to Zeocin, followed by extra-circuit mutations that can shift the entire rtTA response upward, recreating bistability. Filled and open circles denote stable and unstable steady states, respectively. **(D)** Gene expression histograms of evolved population replicates hyperinduced in D6Z0 after the end of experimental evolution. A high expression peak in this condition characterizes the presence of quasi- or fully-functional revertant mutants in the population.

Once again, despite the marked appearance and subsequent gradual disappearance of the high expressing peak in D2Z2, we found no additional rtTA coding sequence mutations in Missense 3 populations. Sanger sequencing from day 14 detected only one intra-circuit mutation in some clones: the deletion of one *tetO2* site upstream from yEGFP::zeoR, while the other *tetO2* site remained intact. Interestingly, all individual Missense 3 clones that acquired a *tetO2* deletion had unimodal gene expression, whereas the clones that lacked this mutation were bistable in D2Z2 at the end (Table S5).

Mathematical modeling (SI model) suggested that gene circuit dysfunction in Missense 3 is due to a drop in rtTA activator capacity (Figure 5C), rather than reduced inducer sensitivity as for Missense 1 and 2. This means that the rtTA synthesis rate curve collapses downward (Figure 5C) instead of shifting rightward in such dysfunctional mutants. Importantly, this collapse implies that the elbow-shaped rtTA loss curve will always be above the collapsed sigmoid during normal growth, so the curves will miss intersecting each other again regardless of the Doxycycline level. Yet, a bistable region can still be reached if slow growth/dilution tilts downward and shifts rightward the rtTA loss curve, causing high-expressors to emerge, despite Missense 3 being completely unresponsive to hyperinduction. Indeed, adding ethanol or Cisplatin but not G418 to D2Z0 indicated that slow growth enables some high expression in Missense 3 (Figure S5). Zeocin in D2Z2 enriches this high expressor fraction further by selection. Mutations elevating the entire sigmoidal synthesis curve could then accelerate growth, thus reestablishing and maintaining bistability (Figure 5C), while also stabilizing the high expressor fraction (SI model). On the other hand, not all mutations elevate expression sufficiently to maintain bistability as growth accelerates, causing the high expressor fraction to diminish in the evolving population, which can be a mixture of such mutation types.

To identify these mutations predicted by the model, we examined WGS data for two replicate populations of Missense 3 (Table S5). Relevant high-frequency mutations in replicate 1 (r1) included 1 bp frameshift deletions in SIF2 and SSN3 as well as a nonsynonymous mutation in SSN2. Missense 3 replicate 3 (r3), on the other hand, carried multiple genomic mutations that spread in the entire population by day 14 of the experimental evolution, including missense mutations in mediator complex components SRB8 and MED6, both of which are involved in RNA polymerase II-dependent transcriptional regulation [44–46] and could elevate both basal and maximal rtTA levels (the entire rtTA synthesis curve) as the model suggested. In addition, a chaperone CCT7 mutation suggests effects on general protein folding [47] and mutations in the SAP185 and EXG1 coding regions suggest altered cell cycle regulation [48, 49] (Dataset S3).

We also hyper-induced the final evolved replicate populations from Missense 3 in D6Z0 to examine PF function at the end of evolution. Surprisingly, two evolved Missense 3 replicates (r1 and r3) had bimodal distributions, indicating that gene circuits in some clones became quasi-functional or fully functional. Missense r2 had a unimodal distribution in D6Z0 indicating that it remained insensitive to hyperinduction (Figure 5D). We observed the same in D8Z0 (Figure S1).

Finally, as for other mutants, we assessed rtTA-dependence of Zeocin resistance in evolved Missense 3 populations and clonal isolates by culturing them in D0Z2. OD600 measurements of original and evolved populations (Figure S9) indicated that evolved Missense 3 replicate populations grew significantly faster than their unevolved ancestor. Interestingly, isolated bimodal Missense 3 clones with mutation-free gene circuits grew slower in D0Z2 than unimodal clones with a *tetO2* site deletion upstream from *yEGFP::zeoR*. This further supports that the *tetO2* site deletion contributes to *rtTA*-independent, but *yEGFP::zeoR*-dependent drug resistance, whereas bimodal Missense 3 clones rely on regained rtTA activity for resistance.

### 2.5. Sorting dysfunctional mutants yields clones with regained rtTA function

So far, a likely common explanation for the observed evolutionary dynamics is the appearance and spread of extra-circuit mutations that accelerate growth of the drug-sensitive, low-expressor subpopulation by elevating *yEGFP::zeoR* basal expression, thereby returning the dynamics towards the unimodal low-expression regime. If this is the case, then separating the high- and low expressors around day 4 should result in different evolutionary dynamics. Since drug selects against ancestral low-expressor cells, the increased selection pressure should cause drug resistance mutations to spread quickly among low-sorted cells, preventing them from generating a substantial high-expressing peak. On the other hand, high-sorted cells would be insensitive to Zeocin and mutations could spread among them only after they generate a low-expressing peak by phenotypic switching. Therefore, high-sorted cells should take longer to become unimodal than low-sorted cells. Also, as discussed above, some quasi- or fully-functional PF revertant clones may exist. If any revertant strains arose, we may be able to isolate them among the high-sorted cells because high expression is the hallmark of PF functionality.

To test these hypotheses, we separated the low-expressor and high-expressor subpopulations by fluorescence-activated cell sorting (FACS) at the end of day 4 of the original evolution experiment and cultured these high- and low-sorted subpopulations separately in the D2Z2 condition for 12 days (Figure 6). As predicted, Missense 1 and 2 low-sorted subpopulations remained in the low-expression state throughout this experiment, indicating rapid takeover by genomic mutations conferring drug resistance. Accordingly, the corresponding high-sorted subpopulations generated bimodal distributions that became unimodal only after a few days as expected.

**Figure 6.**
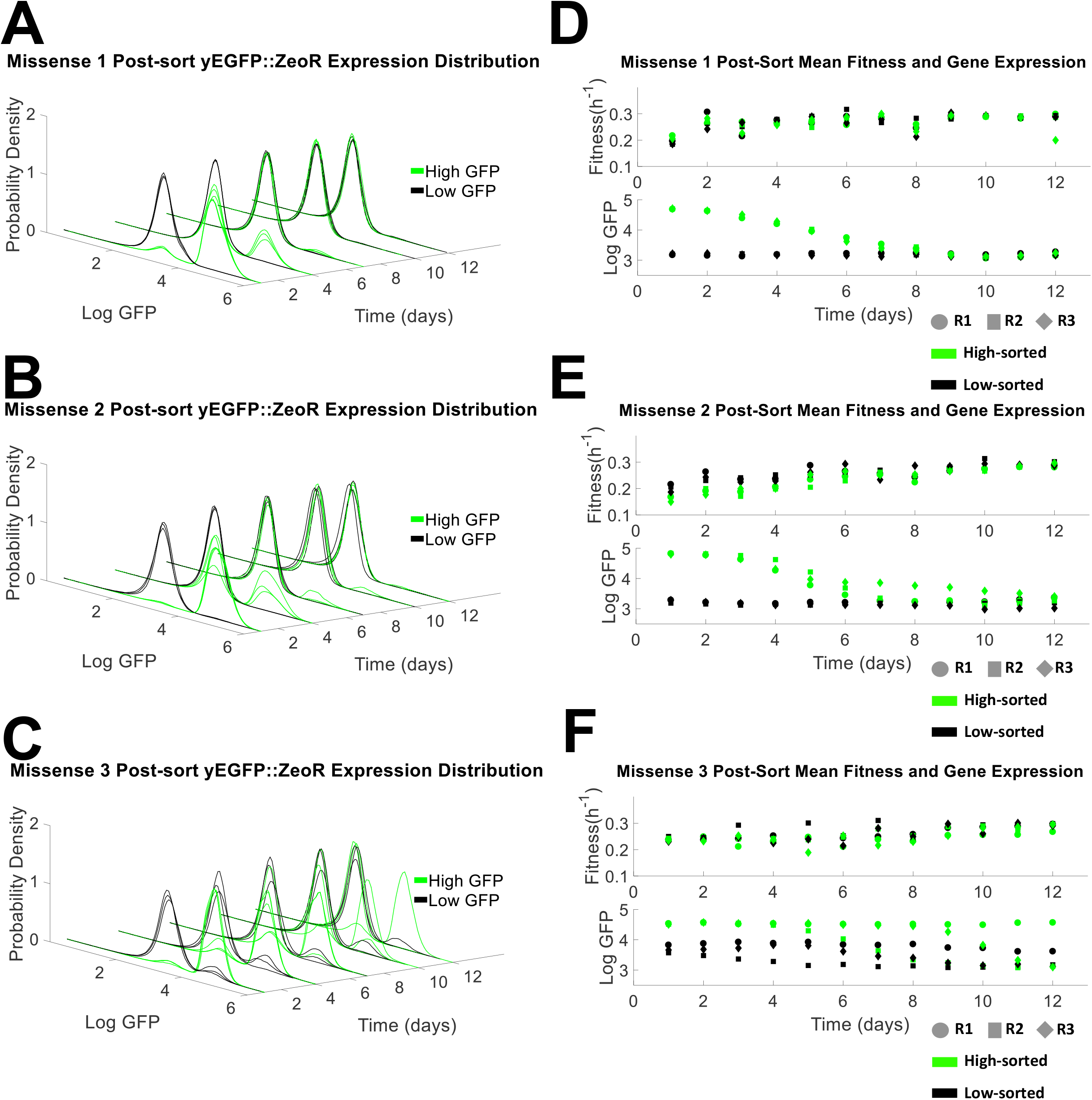
Post-sort evolutionary dynamics of low-sorted and high-sorted subpopulations. **(A)** Histograms of *yEGFP::zeoR* expression for Missense 1 low-sorted (black) and high-expressor (green) subpopulations. Low-sorted subpopulations remain in the low expression state. High-sorted subpopulations give rise to low-expressors within one day after sorting, but then eventually low-expressors dominate. **(B)** Histograms of *yEGFP::zeoR* expression for Missense 2 low-sorted (black) and high-sorted (green) subpopulations. Low-sorted subpopulations remain mostly in the low expression state. High-sorted subpopulations give rise to low-expressors within one day after sorting and eventually low-expressors dominate. **(C)** Histograms of *yEGFP::zeoR* expression for Missense 3 low-sorted (black) and high-sorted (green) subpopulations. For two replicates (r2, r3), low-sorted subpopulations give rise to a high-expressor subpopulation, which then diminishes, while high-sorted subpopulations give rise to low-expressors within one day after sorting and eventually low-expressors dominate similar to Missense 1 and 2. The third replicate (r1) low-sorted subpopulation gives rise to a high-expressor peak and remains bimodal until the end, while the high-sorted subpopulation gives rise to a small low-expressor subpopulation within one day after sorting, but remains bimodal until the end, with a dominant high expressor peak. **(D)** Top: Population fitness of Missense 1 low-sorted (black) and high-sorted (green) subpopulations over the course of 12 days. Bottom: Mean *yEGFP::zeoR* expression of Missense 1 low-sorted (black) and high-sorted (green) subpopulations. **(E)** Top: Population fitness of Missense 2 low-sorted (black) and high-sorted (green) subpopulations over the course of 12 days. Bottom: Mean *yEGFP::zeoR* expression of Missense 2 low-sorted (black) and high-sorted (green) subpopulations. **(F)** Top: Population fitness of Missense 3 low-sorted (black) and high-sorted (green) subpopulations over the course of 12 days. Bottom: Mean *yEGFP::zeoR* expression of Missense 3 low-sorted (black) and high-sorted (green) subpopulations.

As opposed to Missense 1 and 2, Missense 3 low-sorted subpopulations transiently gave rise to new high-expressor subpopulations (Figure 6A-C), indicating either that basal expression-elevating mutants spread much more slowly than in Missense 1 and 2, or that revertant strains arose. For low-sorted Missense 3 replicates r2 and r3, the high-expressor peak gradually diminished within ∼ 9 days. In contrast, low-sorted Missense 3 r1 remained stably bimodal from day 5 throughout the end of this 12-day post-sort experiment.

High-sorted subpopulations from all three Missense 3 replicates gave rise to low-expressor cells within one day after flow sorting. Eventually, the high-expressor peak diminished in all high-sorted subpopulations, except as above, for Missense 3 r1, which remained stably bimodal, biased towards the high expression state for the last 7 days post-sort (Figure 6A-C). Therefore, some Missense 3 r1 clones must be revertant strains with stable high expression.

To identify PF circuit genetic changes underlying these phenotypes, we Sanger-sequenced the *yEGFP::zeoR* and *rtTA* regions of five such individual revertant clones. Once again, the PF sequence in revertant clones was genetically identical to the original Missense 3 mutant. WGS revealed one mutation in the SSN2 gene that spread in the high-sorted Missense 3 r1 population over time (reaching 62.2% on day 12 post-sort), suggesting that it confers a survival benefit and is associated with the stable high expression peak in this population. This mutation was also present at lower frequencies (5.6% and 6.6%) in the respective unsorted and low-sorted populations, further supporting its association with functional reversion. The SSN2 gene encodes an RNA polymerase II mediator complex subunit essential for transcriptional regulation, resembling the general transcriptional-regulatory function of mutations in Missense 2 and Missense 3 r3 (Tables S3, S5). However, the dynamic consequences of such gene expression increases depend on the original mutation – i.e., whether it shrinks or right-shifts the sigmoidal rtTA-synthesis function.

If fully-functional revertant strains arose from Missense 3, they should be clearly bimodal in D2Z0. Thus, to further understand the behavior of revertant strains, we studied the yEGFP::zeoR fluorescence distribution in five individual Missense 3 r1 clones at the end of the post-sort experiment (on day 12) in D2Z2, D0Z2, D2Z0, and D0Z0 for four days (Figure S3). Based on these experiments, Missense 3 r1 revertant clones fell into two phenotypic categories: some had approximately equal peak heights in D2Z2 and D2Z0, whereas other clonal populations consisted almost exclusively of high expressors (Figure 7A,B and Figure S2) like the ancestral PF. Remarkably, the mutation(s) conferred stable high expression even without Zeocin. Additionally, maximum expression shifted leftward, causing the peaks to approach each other compared to the ancestral PF, as predicted by sigmoid elevation in the SI model. Remarkably, the distributions were very similar in D2Z2 and D2Z0, indicating that the revertant strains were drug resistant, so Zeocin did not select for high expression.

**Figure 7.**
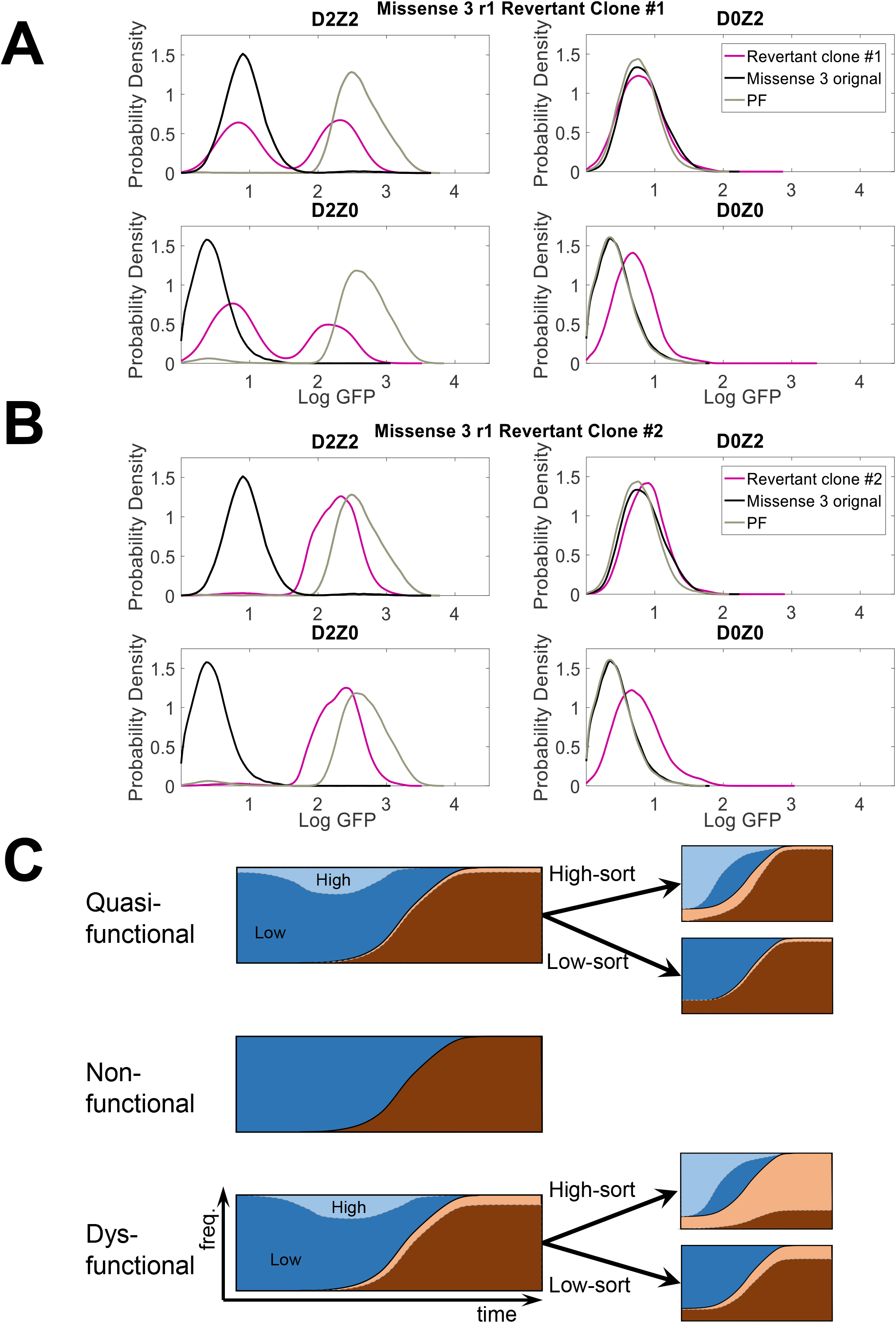
Phenotypic characterization of revertant clones. **(A)** Clone 1 from day 12 of high-sorted Missense 3 r1 populations is a representative of revertant clones with approximately equal gene expression peaks in D2Z2 and D2Z0. **(B)** Clone 2 from day 12 of high-sorted Missense 3 r1 populations is a representative of Missense 3 clones with predominantly high expressors. Histograms were recorded after maintaining each population over 4 days in each indicated condition. **(C)** Muller plots illustrating time-dependent phenotypic and genotypic frequencies (evolutionary dynamics) during pre- and post-sort experimental evolution for quasi-functional, nonfunctional and dysfunctional PF mutants. Different shades indicate identical genotypes, but different phenotypes in bistable populations, the lighter shade corresponding to high expression. Different colors indicate different genotypes. The plots ignore competing genotypes for simplicity. For the dysfunctional mutant Missense 3 the plots illustrate only the revertant clone.

## Discussion

Loss of function is widely observed in experimental evolution studies [5, 8, 13–16], presumably because of the large supply of mutations having these effects. While it is always difficult to extrapolate from the laboratory to natural environments, these observations suggest that loss-of-function may be a common mode of adaptation to a new environment. This raises the question of how populations could regain such lost functions when the environment changes back to a prior state where that function conferred a selective advantage. This question also has great practical significance for synthetic biology, where we may be able to use evolution to resurrect evolutionarily broken synthetic biological systems in medical, environmental or extraterrestrial applications where direct human intervention is difficult or possibly ineffective.

To this end, we investigated evolutionary reversibility by evolving seven yeast strains with broken PF gene circuits, each with a different rtTA mutation, in conditions where regaining gene circuit function would be beneficial. The results revealed various classes of evolutionary dynamics for quasi-functional, dysfunctional, or nonfunctional mutants (Figure 7C). Revertant clones arose only in dysfunctional mutant populations through extra-circuit mutations. After evolving the broken PF mutants, we found no compensatory coding-sequence mutations inside the gene circuits in any populations. Instead, many new extra-circuit mutations increased yEGFP::ZeoR levels, thereby improving drug resistance, without restoring broken PF function, indicating that evolution adopts alternate paths if they are available [50, 51]. Restricting such alternate paths (e.g., by using higher drug concentrations or by preventing extra-circuit genomic mutations [52]) may facilitate functional reversions in future experiments. Importantly, some extra-circuit mutations did re-enable PF gene circuit function by elevating rtTA expression, indicating that evolutionary reversion is possible without any new mutations in rtTA or even the gene circuit. This is broadly consistent with another recent study that evolved *E. coli* with loss of function mutations in a particular enzyme, showing some direct revertant mutations in the functional locus but also indirect adaptations elsewhere in the genome [53].

We discovered interesting interactions between intracellular gene circuit dynamics, population dynamics and evolutionary dynamics [54]. Specifically, slow growth rate was not only a fitness parameter, but also a potential enabler of dynamic shifts leading to beneficial early phenotypic and later genetic changes. This is reminiscent of the emerging concept of growth rate as a global regulator of the cell’s transcriptional state [55, 56]. The potential implications are remarkable, since a slow-growing dysfunctional mutant can rapidly develop a large high expression peak, appearing to be reverted. However, these early expression shifts are not truly evolutionary reversions – they simply result from gaining access to a bistable regime that existed for the entire time but was inaccessible through hyperinduction. Indeed, other conditions that decelerate growth similarly to Zeocin, ethanol and Cisplatin could also boost bistability for such mutants. To combat drug resistance, it will be important to understand how often similar growth-related dynamic shifts [38, 39, 57] can happen in natural networks of bacteria, yeasts, and mammalian cells. Furthermore, the antibiotic’s mechanism of action matters for survival and evolution. As the mathematical model suggested stressors that reduce growth without directly interfering with protein synthesis enabled access to a bistable regime. On the other hand, antibiotics such as G418 that directly inhibit translation [58], might prevent protein expression-dependent dynamical shifts, and consequently the emergence of high-expressors. Future studies evaluating the effect of different antibiotic mechanisms of action on gene network dynamics and evolution will be important and informative.

Sanger sequencing revealed only one intra-circuit mutation across all different tested mutants: a 42bp deletion eliminating the first *tetO2* operator site upstream of the *yEGFP::zeoR* coding region. The other *tetO2* site upstream from *yEGFP::zeoR* and the two *tetO2* sites upstream from rtTA remained intact, leaving positive feedback unaltered while still enabling *yEGFP::zeoR* activation by rtTA. We could not link the observed phenotypes and the identified *tetO2* site deletion specifically, except for Missense 3 where one *tetO2* site was absent only in monostable populations, while bistable clones had intact gene circuits. Notably, *tetO2* site deletions occurred in nonfunctional mutants and also in cells previously evolved in D0Z2 [9], suggesting that these deletions contribute to elevating basal yEGFP::zeoR expression independently of rtTA. Establishing the functional and dynamic effects of single versus double *tetO2* sites in front of *yEGFP::zeoR* requires further investigation. Extra-circuit mutations in various genes such as *SRB8*, *MED6*, *CCT7*, *NUP159*, and others suggest alterations in the transcription and protein folding processes, which will require further studies.

Upon sorting and separately culturing low- and high-expressor subpopulations from evolving populations with quasi-functional and dysfunctional mutants, only the latter contained revertant clones that reestablished and maintained bistability. Hence, we successfully generated fully revertant clones capable of stably reactivating the mutant PF circuit. Interestingly, the reversion could be linked to an extra-circuit mutation in the *SSN2* gene, pointing to functional alterations in the RNA polymerase II-mediator complex controlling transcriptional regulation, and consequently conferring drug resistance through rtTA and yEGFP::zeoR protein levels. Compared to the ancestral PF and the original Missense 3 gene circuit, the revertant strains were Zeocin-resistant (Figure S10) yet high expression was not costly, indicating that breaking and then recovering the gene circuit function resulted in PF cell lines with functional gene circuits that are more robust to evolution and environmental perturbations.

Fusing an antibiotic resistance cassette to genes of interest would likely be useful to restore the function of other activator-based gene circuits. The regulators of gene circuits employing only repressors (such as the toggle switch) would be less likely to degrade by evolution in eukaryotes where steric repressors tend to be less toxic than activators, so the evolutionary pressure to mutate them would be lower. Nonetheless, fusing an antibiotic resistance gene to target genes and applying drug selection should allow regain of function even for repressor-based gene circuits if they break down. Moreover, auxotrophic positive- and negative-selection markers (such as URA3) could be similarly employed as handles for evolutionary restoration, with the advantage that selection could be applied both for low and high expression in yeast.

As opposed to our method, directed evolution studies in the past optimized enzymes and metabolic pathways [33, 34] to avoid the difficulties of rational troubleshooting and screening. Such studies focused on individual proteins, mutagenizing coding sequences and screening for optimal variants. In particular, a study titled “Directed evolution of a genetic circuit” in fact mutagenized the coding sequence of a single transcription factor and then screened for optimal variants without actually performing network evolution [33], that is, allowing the entire network and host genome to change naturally. Other studies investigated evolutionary degradation of gene circuits but did not seek to recover or improve gene circuit function by experimental evolution [9, 30].

To conclude, our results highlight the versatility of drug resistance mechanisms, including dynamical consequences of slow growth, and exemplify how yeast and possibly mammalian [23] gene circuit evolution can reveal interesting dynamical and biological behaviors, such as host-genomic mutations repairing lost gene circuit function.

## Materials and Methods

### Yeast strains

We used previously evolved [9] YPH500 haploid *Saccharomyces cerevisiae* clones (α, ura3-52, lys2-801, ade2-101, trp1Δ63, his3Δ200, leu2Δ1; Stratagene) with wild-type and seven rtTA-mutant PF synthetic gene circuits [6] stably integrated into chromosome XV near the *HIS3* locus (Table S1). A 2% weight of galactose Synthetic Dropout (SD) culture medium with appropriate supplements (-his, -trp) was used to maintain auxotrophic selection (all reagents from Sigma). Cells were grown in SD-his-trp + 2% galactose plus Doxycycline and Zeocin as indicated at 30°C, shaking at 300 rpm (LabNet 311DS shaking incubator).

### Flow-cytometry

The BD FACSAria™ III at the Stony Brook Flow-cytometry facility was used for cell sorting.

The BD Accuri™ C6 flow-cytometer was used to collect data for hyperinduction, experimental evolution for sorted and unsorted populations.

The BD FACSCalibur™ at the Stony Brook Flow-cytometry facility was used to collect data for slow growth rate and phenotyping experiments.

### Estimation of population fitness

We counted cells daily using the Nexcelom Bioscience Cellometer Vision cell counter. The daily (*T*=24h) resuspension of a low number of cells (10^5^) ensures that cells remain in the exponential growth phase throughout these experiments. Therefore, based on Nexcelom cell counts before each resuspension, we can use the exponential growth equation to estimate the population growth rate, 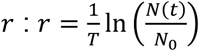.

### Hyperinduction and slow growth rate experiments

To classify quasi-functional, dysfunctional and nonfunctional rtTA mutants, we cultured them in 6 µg/mL (D6Z0) or 8 µg/mL (D8Z0) Doxycycline (Thermo Fisher Scientific) added to SD-his-trp for 24h. Flow-cytometry (BD Accuri™) was performed to measure gene expression levels.

To assess the effect of slow growth rate on mutant and ancestral PF circuit dynamics, we cultured mutant (Missense 1, 2, 3, Nonsense and Deletion) and ancestral PF strains in 2 µg/mL Doxycycline (D2Z0) +7.5% 200 proof ethanol (Pharmco) for 4 days. Cells were resuspended at a 10^5^ cells/mL density every day. We measured cell counts daily and protein expression by flow-cytometry on days 1, 2 and 4 (BD FACSCalibur™). Additionally, we conducted the same experiment using Cisplatin (Selleckchem) at a concentration of 40ug/mL, instead of ethanol.

### Experimental evolution

Previously [9], mutant strains were isolated and stored in 27% glycerol (Thermo Fisher Scientific) at −80C. Seven mutant strains were picked from the −80C stock, streaked onto SD -his -trp 2% glucose plates (all reagents from Sigma) and incubated at 30°C for 2 days. One isolated colony from each plate was cultured in liquid SD -his -trp 2% galactose media at 30°C & 300rpm for 24 hours. For each strain, 10^5^ cells were resuspended into the D2Z0 (2 µg/mL Doxycycline) and D2Z2 (2 µg/mL Doxycycline (Thermo Fisher Scientific), 2mg/mL Zeocin (Thermo Fisher Scientific) conditions (3 replicates each) every ∼24h. Fluorescence (BD Accuri™) and cell counts were measured daily. Each day, samples were stored in 27% glycerol at −80C. The experiment lasted for 14 days.

### Zeocin only growth curve monitoring

Original mutants and evolved replicates were cultured in the D0Z2 (2mg/mL Zeocin (Thermo Fisher Scientific)) condition at a concentration of 10^5^ cells/mL. Growth curves were monitored through OD600 measurements (every 20 minutes for 24 hours) using the Tecan Infinite® 200 PRO plate reader. The following strains were monitored in D0Z2:

- Original (unevolved) ancestor strains of each mutant.
- Replicate 3 (WGS replicate) of evolved populations of each mutant, from day 14 of experimental evolution, except for Missense 3 where all replicates were tested.
- Sequenced and phenotyped clonal isolates (shown in Tables S3-S5) for each mutant.

In addition, for control and comparison purposes, we monitored the growth of ancestral PF in D0Z0, D2Z2 and D0Z2.

### Cell Sorting

Missense 1, 2 and 3 populations were FACS-sorted at day 4 of the experiment at the Stony Brook flow cytometry facility using the BD FACSAria™ III. Sorting parameters were set on the BD FACSDiva™ 8.0 software. High-sorted cells were collected from the highest 5-10% of the high-expressor subpopulations. Collected low- and high-expressor subpopulations were cultured separately in D2Z2 for 12 days. Flow-cytometry (BD Accuri™) and cell counts were performed daily. Each day, samples were stored in 27% glycerol at −80C.

### DNA isolation

DNA was extracted from isolated single clones for Sanger sequencing or from evolved populations for WGS using the MasterPure™ Yeast DNA Purification Kit (Lucigen) as per manufacturer’s instructions.

### Sanger sequencing

Sanger sequencing was performed on 10 isolated evolved clones from each unsorted mutant population at day 14 and 5 isolated clones from the sorted revertant Missense 3 replicate at day 12. For primers, see Table S2. Samples were sequenced at the DNA Sequencing Facility at Stony Brook University using the BigDye Terminator v3.1 sequencing kit, BigDye XTerminator Purification kit and the 3730 DNA Analyzer (all from Applied Biosystems). Results were analyzed using the SnapGene software sequence alignment tool.

### Whole genome sequencing (WGS) analysis

WGS was performed on Missense 1 r3, Missense 2 r3 and Missense 3 r1 and r3 at days 1, 3 and 14 for unsorted populations and days 1, 5 and 12 for the sorted populations. WGS was also performed on r3 of each of the other mutants on day 14 only. Samples were sent to Novogene Co., Ltd. where library preparation was performed using the NEBNext® Ultra II kit (New England Biolabs) and sequencing was performed by Illumina HiSeq-4000 at a 165x coverage.

To analyze the sequencing data, we used the S288C genome (RefSeq accession numbers NC_001133.9, NC_001134.8, NC_001135.5, NC_001136.10, NC_001137.3, NC_001138.5, NC_001139.9, NC_001140.6, NC_001141.2, NC_001142.9, NC_001143.9, NC_001144.5, NC_001145.3, NC_001146.8, NC_001147.6, NC_001148.4, NC_001224.1) as a reference with the PF circuit inserted computationally on chromosome 15 between the MRM1 and HIS3 genes. We processed the reads and this modified reference genome using the breseq 0.33.1 pipeline on default settings for polymorphism prediction [59]. The pipeline produced lists of inferred variants (including SNPs, insertions, and deletions) and their estimated frequencies in the population for each sequencing sample. Subsequently, inferred variants were filtered down according to the following three criteria: (i). Variant cannot appear in samples from different populations (e.g., Missense 1 and Missense 3). (ii).Variant must appear in multiple samples if it appears in a population with multiple samples. (iii).Variant must appear >10% in at least one sample.

Here we highlight mutations as potentially causal if they increase in frequency over time or if they are absent at the beginning but are detected in high frequencies (>50%) at the end of the evolution experiment. All detected variants are reported (Datasets S1-S7).

### Phenotyping of isolated clones

To understand the effect of the *tetO2* deletion detected upstream of *yEGFP::zeoR* by Sanger sequencing on PF dynamics, one clone from each possible PF genotype found (presence of *tetO2* or deletion of *tetO2*) was cultured in D2Z2 for 24h. Flow-cytometry was performed to obtain a protein expression distribution (BD FACSCalibur™).

To compare PF to the revertant Missense 3 r1 mutant, PF and the five Sanger-sequenced revertant clones were cultured in D0Z0, D2Z0, D2Z2 and D0Z2 for 4 days. Flow-cytometry was performed to obtain a protein expression distribution on each day (BD FACSCalibur™).

### Quantitative modeling and software

Flow cytometry data were gated based on forward and side scatter using FCS Express and then exported for subsequent analysis. All experimental data were analyzed and plotted in MATLAB R2016b. WGS data were analyzed using the breseq 0.33.1 software. Ordinary differential equation (ODE) models were developed, then their steady states were analyzed on paper and by custom-written scripts using the function *roots* in MATLAB R2018a. See the SI model for details.

## Acknowledgments

We would like to thank Rebecca C. Connor and Todd Rueb for their help in performing the FACS sorting and the flow-cytometry readings at the Stony Brook flow cytometry facility. We also thank Oleksandra Romanyshyn and Michael Tyler Guinn for facilitating the timely execution of experiments. Finally, we would like to thank all Balázsi lab members for helpful discussions and feedback. This work was funded by the NIH/NIGMS MIRA grant R35 GM122561 (GB), and by the Laufer Center for Physical and Quantitative Biology (GB), and the Swiss National Science Foundation Ambizione grant PZ00P3_180147 (MM).

## Author contributions

G.B. and M.K.G. designed the study. M.K.G. executed all experiments. M.K.G. analyzed flow-cytometry, growth rate and Sanger sequencing data. M.M. performed the whole genome sequencing analysis, G.B. designed the mathematical model and corresponding computational framework. G.B. and M.K.G. interpreted the data, and G.B. and M.K.G. wrote the manuscript.

