## Supplementary Information for "Evolutionary regain of lost gene circuit function"

### 1. Supporting Tables and Datasets

| Mutant | Position in rtTA | Position in S288C+PF chr15 | Nucleotide effect | Amino acid effect | Phenotypic effect | Evolved population: Gonzalez et al., 2015 |
|---|---|---|---|---|---|---|
| **Missense 1** | 189 | 729,794 | C→G | His→Gln | Quasi-functional | D2Z0-12hr-r2 |
| **Missense 2** | 562 | 730,167 | T→C | Phe→Leu | Quasi-functional | D2Z0-24hr-r2 |
| **Missense 3** | 275 | 729,880 | G→A | Ser→Asn | Dysfunctional | D2Z0-24hr-r3 |
| **Missense 4** | 13 | 729,618 | G→T | Asp→Tyr | Nonfunctional | D2Z0-24hr-r3 |
| **Nonsense** | 442 | 730,047 | G→T | Glu→Stop | Nonfunctional | D2Z0-24hr-r2 |
| **Deletion** | 651 | 730,256 | 78bp deletion | 26 amino acid deletion | Nonfunctional | D2Z0-24hr-r3 |
| **Duplication** | 95 | 729,700 | 30bp duplication | 10 amino acid duplication | Nonfunctional | D2Z0-24hr-r1 |

**Table S1**. Genotypes of the 7 mutants before the evolution experiment.

| Primer name | Sequence |
|---|---|
| TRP-f | ATGTCTGTTATTAATTTCACAGGTAGTTC |
| rtTA-seq-int-r-cg | CGACTTGATGCTCTTGTTCTTCCAATACGCAACC |
| rtTA-seq-int-f-cg | GCCAACAAGGTTTTTCACTAGAGAATGCATTATATG |
| Tetreg-AflII-f | GCGCCTTAAGGCGCCACTTCTAAATAAGCGAATTTC |
| rtTABamHI2-f | GCGCGGATCCATGTCTAGATTAGATAAAAGTAAAG |
| FFF-XhoI-r | GCGCCTCGAGTTAACCTGGCAACATATCTAAATCAAAGTCATC |
| Backbone-r | CGCGTTGGCCGATTCATTAATGC |
| His-f | ATGACAGAGCAGAAAGCCCTAGTAAAGC |
| ZeoR-XhoI-r | GCGCCTCGAGTCAGTCCTGCTCCTC |

**Table S2.** Sanger sequencing primers for the PF gene circuit region.

| Mutant | Genomic mutations | | Frequency of genomic mutations at day 14 | *tetO2* deletion in 10 clones | Single clone phenotype in D2Z2 (24h) |
|---|---|---|---|---|---|
| Missense 1 r3 | Phospholipid biosynthesis | *PAH1*<br>T479P (ACG→CCG)<br>(Thr→Pro) | 14.3% | 10/10 | 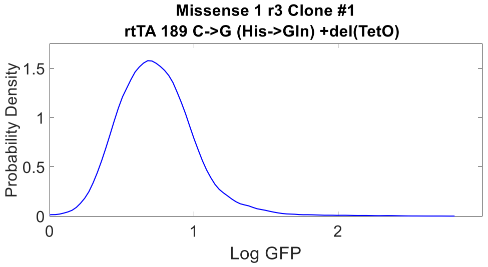 |
| | Histone-binding | *SET3*<br>coding (440/2256 nt)<br>Δ1 bp | 14.5% | | |
| | | *SET3*<br>G147S (GGC→AGC)<br>(Gly→Ser) | 14.4% | | |
| Missense 2 r3 | Mitochondrial metabolism | *Phb2*<br>Y138N (TAC→AAC)<br>(Tyr→Asn) | 100% | 0/10 | 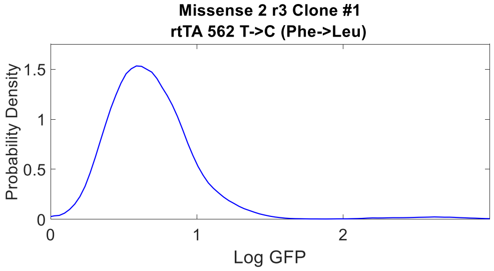 |
| | | *Mdm32*<br>S142C (AGC→TGC)<br>(Ser→Cys) | 100% | | |
| | | *COX1*<br>Intragenic<br>(+762/-125)<br>Δ1bp | 100% | | |
| | Transcriptional regulation | *NUP159*<br>A615A (GCA→GCC)<br>Synonymous Ala | 100% | | |

**Table S3.** Genotyping and phenotyping of individual clones of quasi-functional Missense 1 and Missense 2 mutants. Genomic mutations reported here arose during the evolutionary experiment.

| Mutant | *tetO2* deletion in 10 clones | Single clone phenotype in D2Z2 (24h) |
|---|---|---|
| Missense 4 r3 | 1/10 | 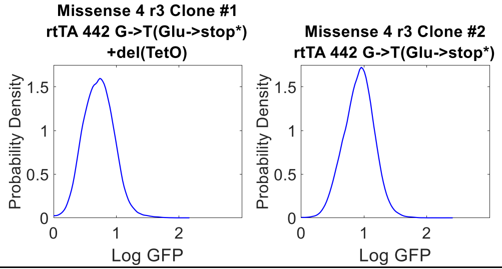 |
| Nonsense r3 | 0/10 | 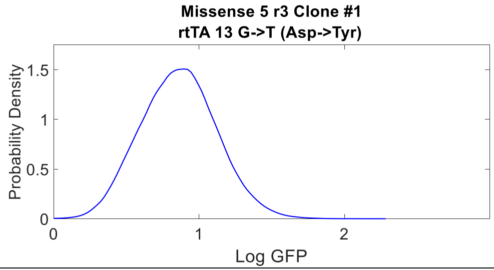 |
| Deletion r3 | 8/10 | 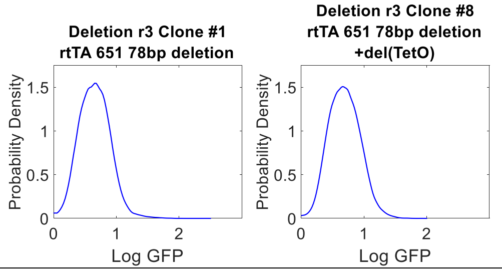 |
| Duplication r3 | 2/10 | 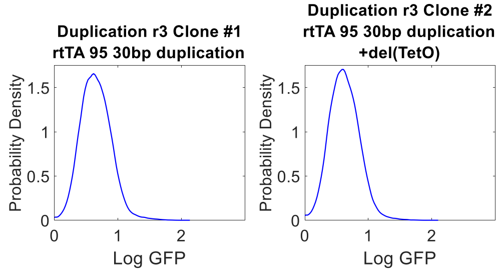 |

**Table S4.** Genotyping and phenotyping of individual clones of nonfunctional Missense 4, Nonsense, Deletion and Duplication mutants. Mutations reported here arose during the evolutionary experiment.

| Mutant | Genomic mutations | | Frequency of genomic mutations at day 14 | *tetO2* deletion in 10 clones | Single clone phenotype in D2Z2 (24h) |
|---|---|---|---|---|---|
| Missense 3 r1 | Chromatin regulation | *SIF2*<br>coding (144/1608 nt)<br>Δ1 bp | 32.6% | 4/10 | 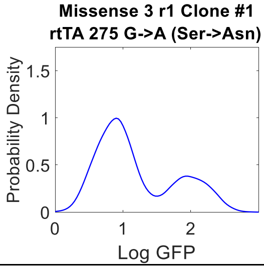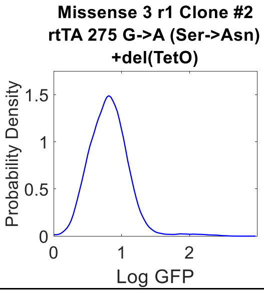 |
| | Transcriptional regulation | *SSN2*<br>V239D (GTT→GAT)<br>Val→Asp | 5.6% | | |
| | Transcriptional regulation | *SSN3*<br>coding (47/1668 nt)<br>Δ1 bp | 34.4% | | |
| Missense 3 r2 | Not sequenced by WGS | | | 10/10 | 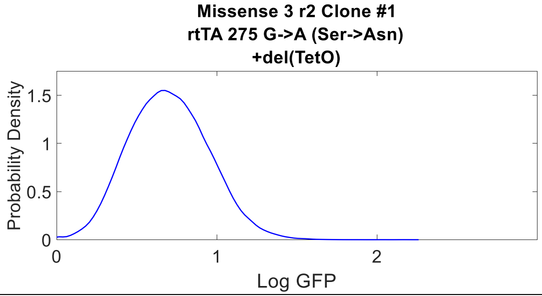 |
| Missense 3 r3 | Transcriptional regulation | *SRB8*<br>P116R (CCT→CGT)<br>(Pro →Arg) | 100% | 0/10 | 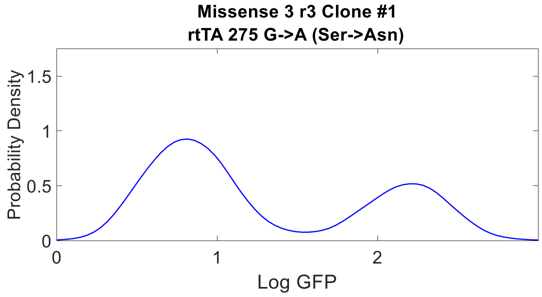 |
| | | *MED6*<br>Y100F (TAT→TTT)<br>(Tyr→Phe) | 100% | | |
| | Protein folding | *CCT7*<br>R469S (AGA→AGT) | 100% | | |
| | Cell Cycle | *SAP185*<br>coding (423/3177 nt)<br>Δ1bp | 100% | | |
| | | *ECM38* →/→ *EXG1*<br>intergenic<br>(+634/-270)<br>(T→A) | 100% | | |

**Table S5.** Genotyping and phenotyping of individual clones of dysfunctional Missense 3 mutant. Genomic mutations reported here arose during the evolutionary experiment.

**List of Whole Genome Sequencing Datasets:**

**Dataset S1.** WGS variants for Missense 1, replicate 3.

**Dataset S2.** WGS variants for Missense 2, replicate 3.

**Dataset S3.** WGS variants for Missense 3, replicates 1 and 3.

**Dataset S4.** WGS variants for Missense 4, replicate 3.

**Dataset S5.** WGS variants for Nonsense, replicate 3.

**Dataset S6.** WGS variants for Deletion, replicate 3.

**Dataset S7.** WGS variants for Duplication, replicate 3.

### 2. Supporting Figures

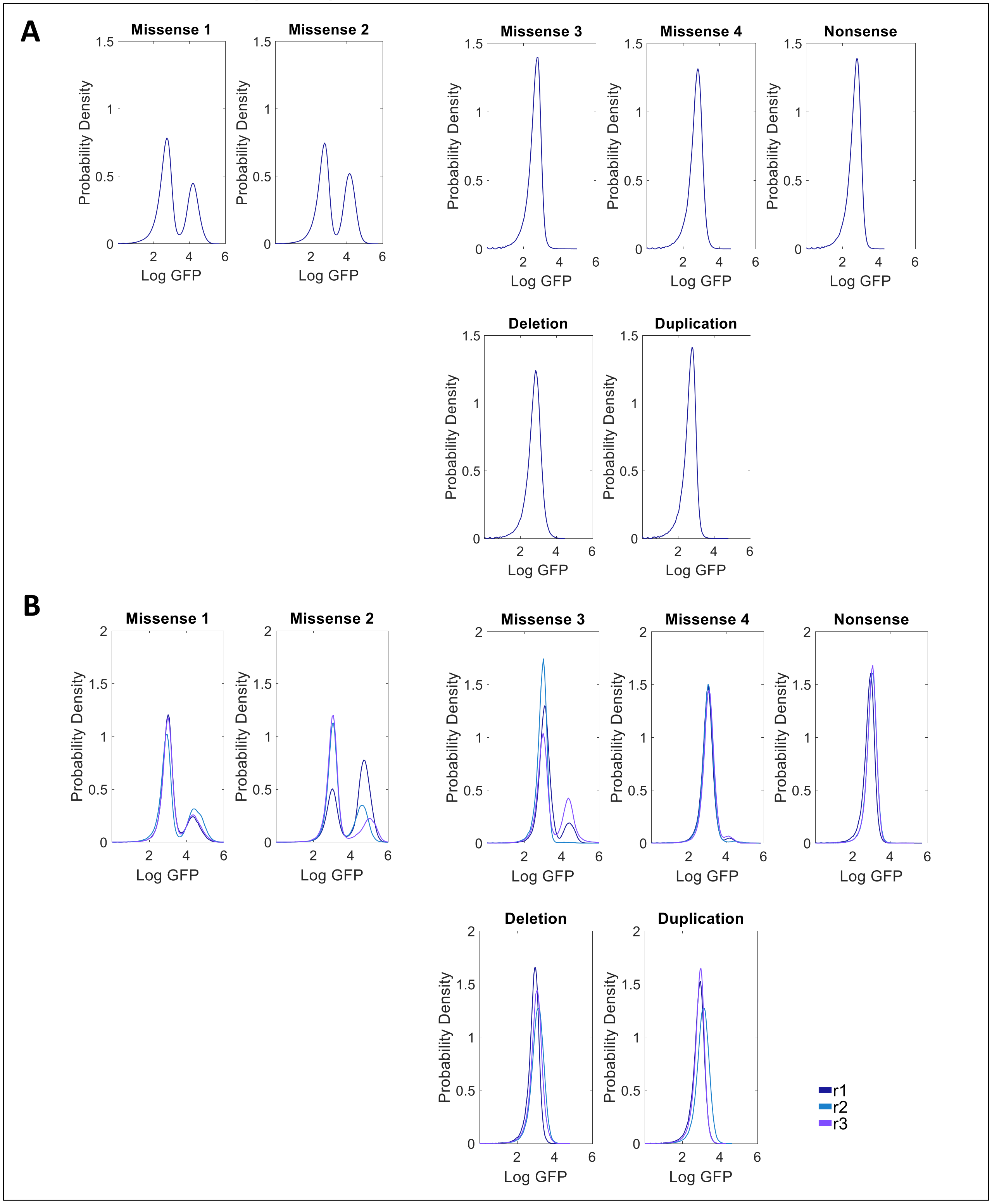

**Figure S1.** (A) Hyper-induction of original mutants in D8Z0 prior to the evolution experiment. (B) Hyper-induction of evolved mutants in D8Z0 following the evolution experiment.

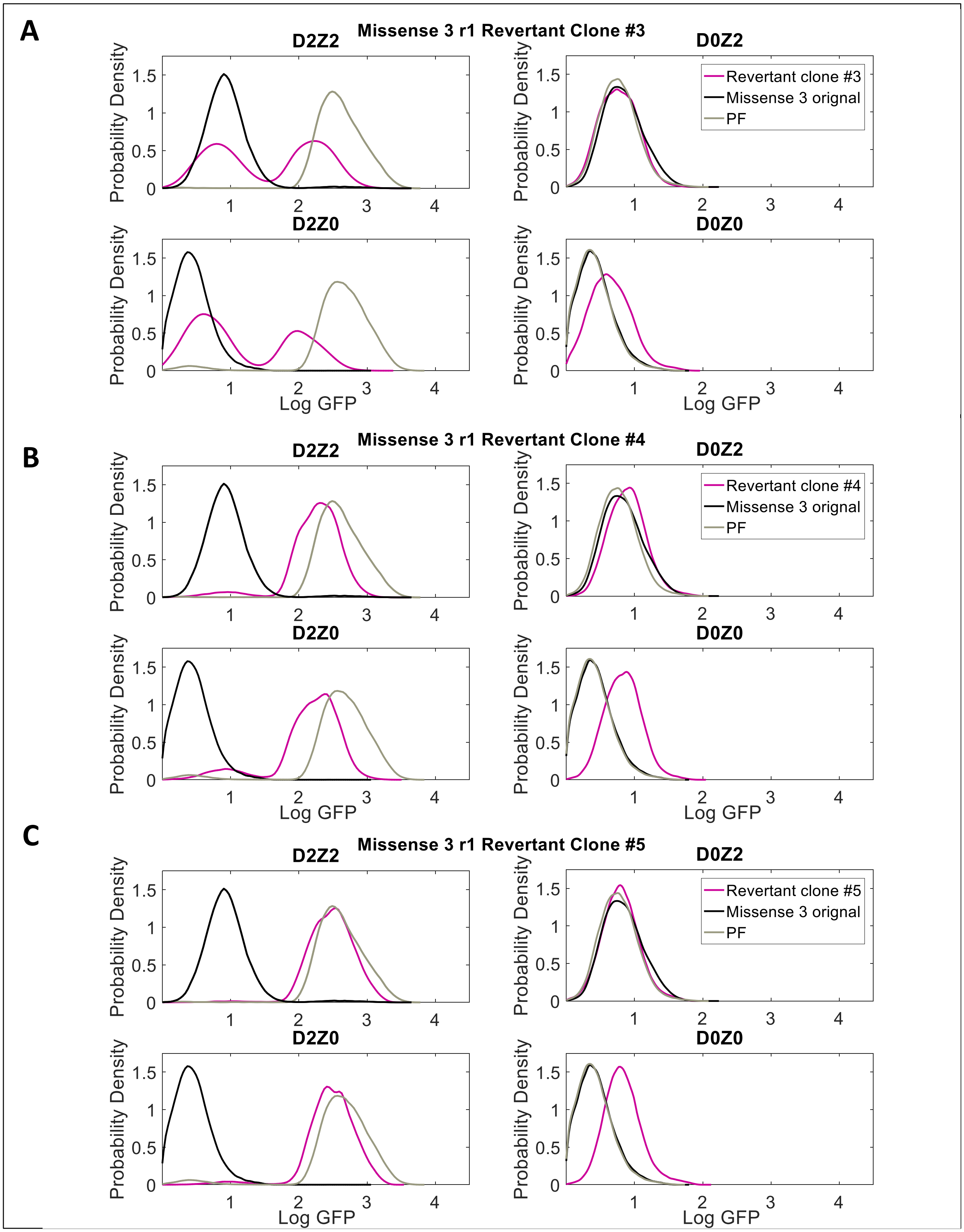

**Figure S2.** Gene expression distributions of three individual clones from revertant Missense 3 replicate r1 in D0Z0, D2Z0, D2Z2 and D0Z2 at day 4.

**A**

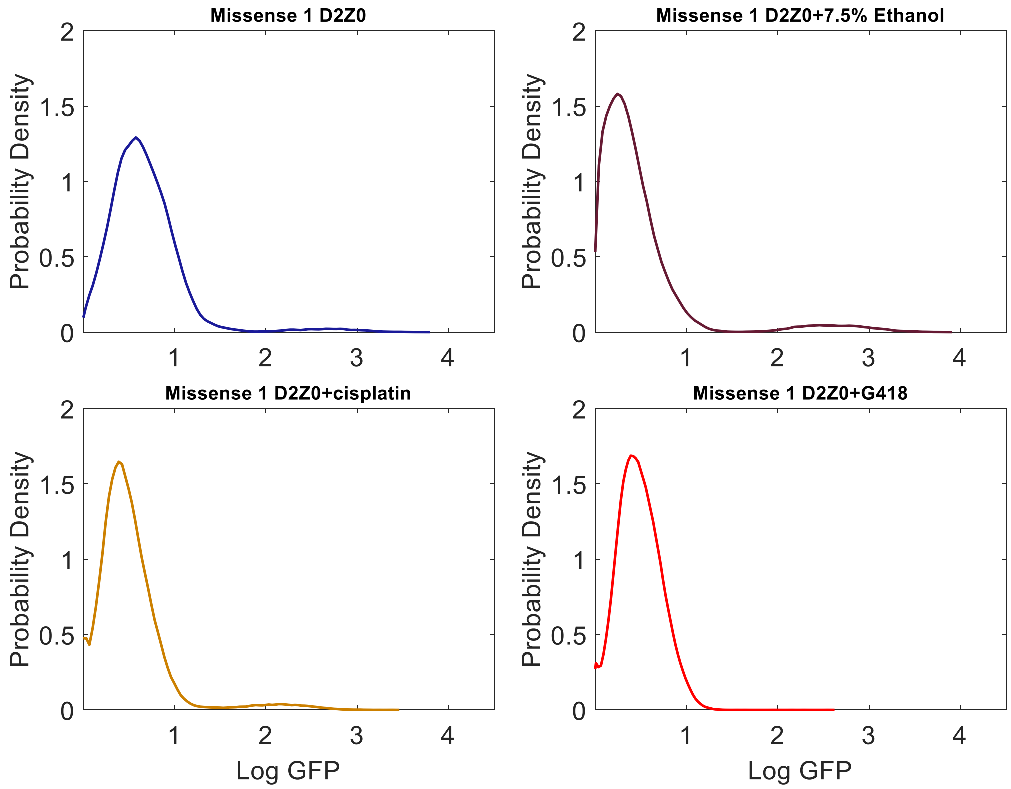

**B**

| Missense 1 D2Z0 g=0.284 | | | |
|---|---|---|---|
| Subpop. | # of events | Percentage | Geometric mean |
| High GFP | 2033 | 1.85% | 198.97 |
| Low GFP | 107754 | 98.14% | 3.42 |

| Missense 1 D2Z0 + 7.5% EtOH g=0.098 | | | |
|---|---|---|---|
| Subpop. | # of events | Percentage | Geometric mean |
| High GFP | 4437 | 4.06% | 185.85 |
| Low GFP | 104935 | 95.92% | 1.89 |

| Missense 1 D2Z0 + Cisplatin (40ug/mL) g=0.093 | | | |
|---|---|---|---|
| Subpop. | # of events | Percentage | Geometric mean |
| High GFP | 3563 | 3.24% | 137.89 |
| Low GFP | 106427 | 96.76% | 2.88 |

| Missense 1 D2Z0 + G418 (40ug/mL) g=0.093 | | | |
|---|---|---|---|
| Subpop. | # of events | Percentage | Geometric mean |
| High GFP | 4 | 0.00% | 171.54 |
| Low GFP | 109986 | 100.00% | 3.05 |

**Figure S3.** Effect of slow growth rate due to ethanol and Cisplatin on Missense 1 distributions in D2Z0 at day 4. (A) Gene expression distributions. (B) Histogram statistics.

**A**

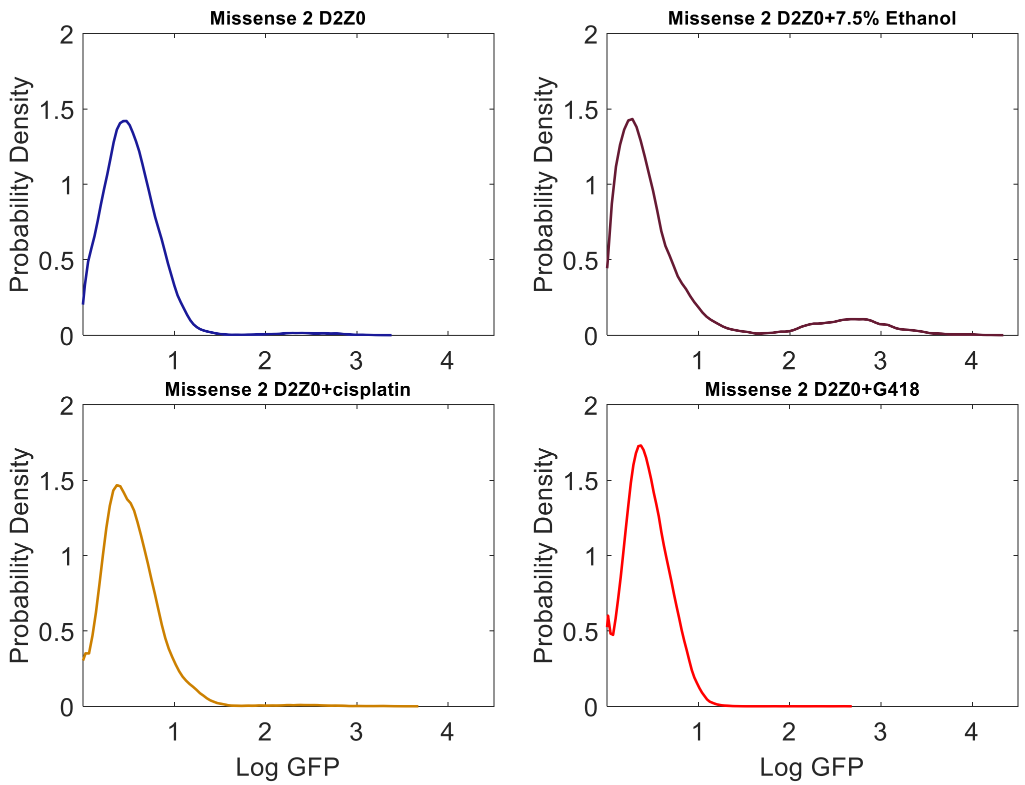

**B**

| Subpop. | # of events | Percentage | Geometric mean |
|---|---|---|---|
| **Missense 2 D2Z0 g=0.269** | | | |
| **Subpop.** | **# of events** | **Percentage** | **Geometric mean** |
| High GFP | 1204 | 1.10% | 131.32 |
| Low GFP | 108599 | 98.90% | 2.67 |
| **Missense 2 D2Z0 + 7.5% EtOH g=0.052** | | | |
| **Subpop.** | **# of events** | **Percentage** | **Geometric mean** |
| High GFP | 2503 | 9.97% | 217.84 |
| Low GFP | 22578 | 89.94% | 2.09 |
| **Missense 2 D2Z0 + Cisplatin (40ug/mL) g=0.127** | | | |
| **Subpop.** | **# of events** | **Percentage** | **Geometric mean** |
| High GFP | 803 | 0.73% | 261.61 |
| Low GFP | 108746 | 99.27 | 3.30 |
| **Missense 2 D2Z0 + G418 (40ug/mL) g=0.138** | | | |
| **Subpop.** | **# of events** | **Percentage** | **Geometric mean** |
| High GFP | 8 | 0.01% | 181.26 |
| Low GFP | 109880 | 99.99% | 2.74 |

**Figure S4.** Effect of slow growth rate due to ethanol and Cisplatin on Missense 2 distributions in D2Z0 at day 4. (A) Gene expression distributions. (B) Histogram statistics.

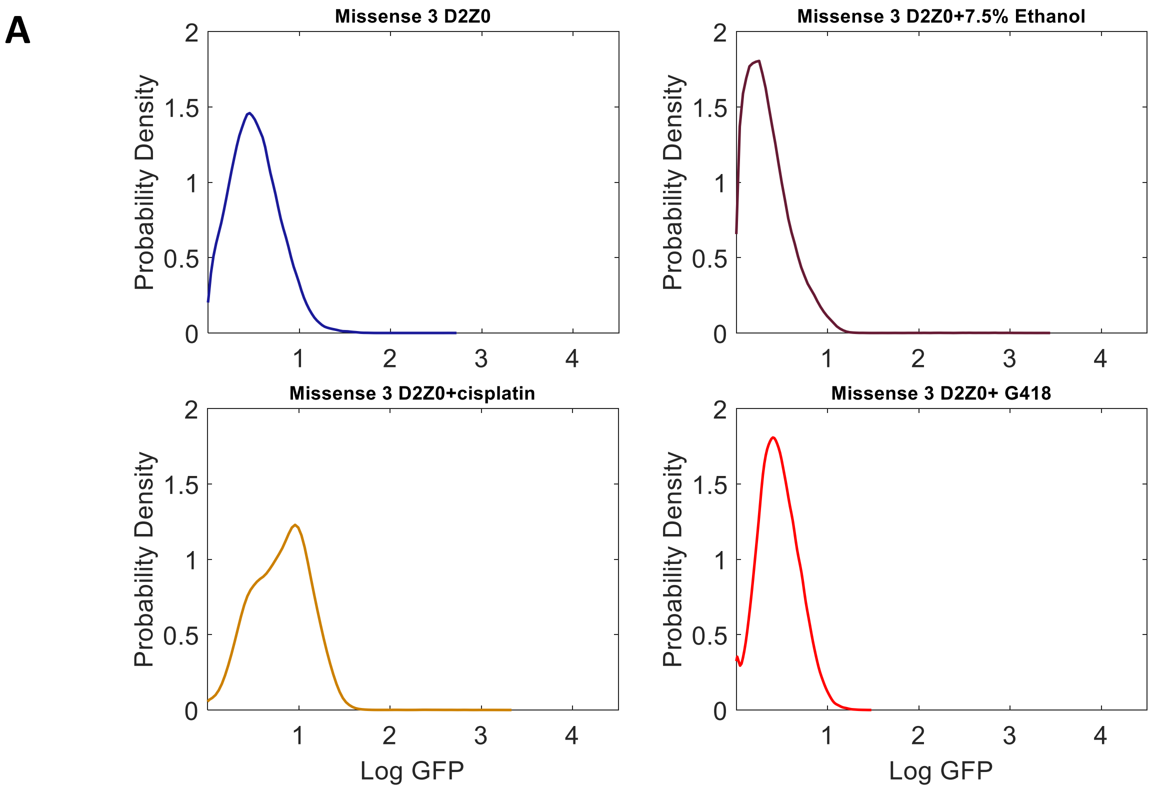

**B**

| Missense 3 D2Z0 g=0.266 | | | |
|---|---|---|---|
| **Subpop.** | **# of events** | **Percentage** | **Geometric mean** |
| **High GFP** | **0** | **0.00%** | **N/A** |
| **Low GFP** | **109892** | **100.00%** | **2.65** |
| **Missense 3 D2Z0 + 7.5% EtOH g=0.048** | | | |
| **Subpop.** | **# of events** | **Percentage** | **Geometric mean** |
| **High GFP** | **74** | **0.07%** | **157.30** |
| **Low GFP** | **108200** | **99.93%** | **1.77** |
| **Missense 3 D2Z0 + Cisplatin (40ug/mL) g=0.079** | | | |
| **Subpop.** | **# of events** | **Percentage** | **Geometric mean** |
| **High GFP** | **52** | **0.09%** | **222.44** |
| **Low GFP** | **57908** | **99.91%** | **6.31** |
| **Missense 3 D2Z0 + G418 (40ug/mL) g=0.115** | | | |
| **Subpop.** | **# of events** | **Percentage** | **Geometric mean** |
| **High GFP** | **0** | **0.00%** | **N/A** |
| **Low GFP** | **109958** | **100.00%** | **2.89** |

**Figure S5.** Effect of slow growth rate due to ethanol and Cisplatin on Missense 3 distributions in D2Z0 at day 4. (A) Gene expression distributions. (B) Histogram statistics.

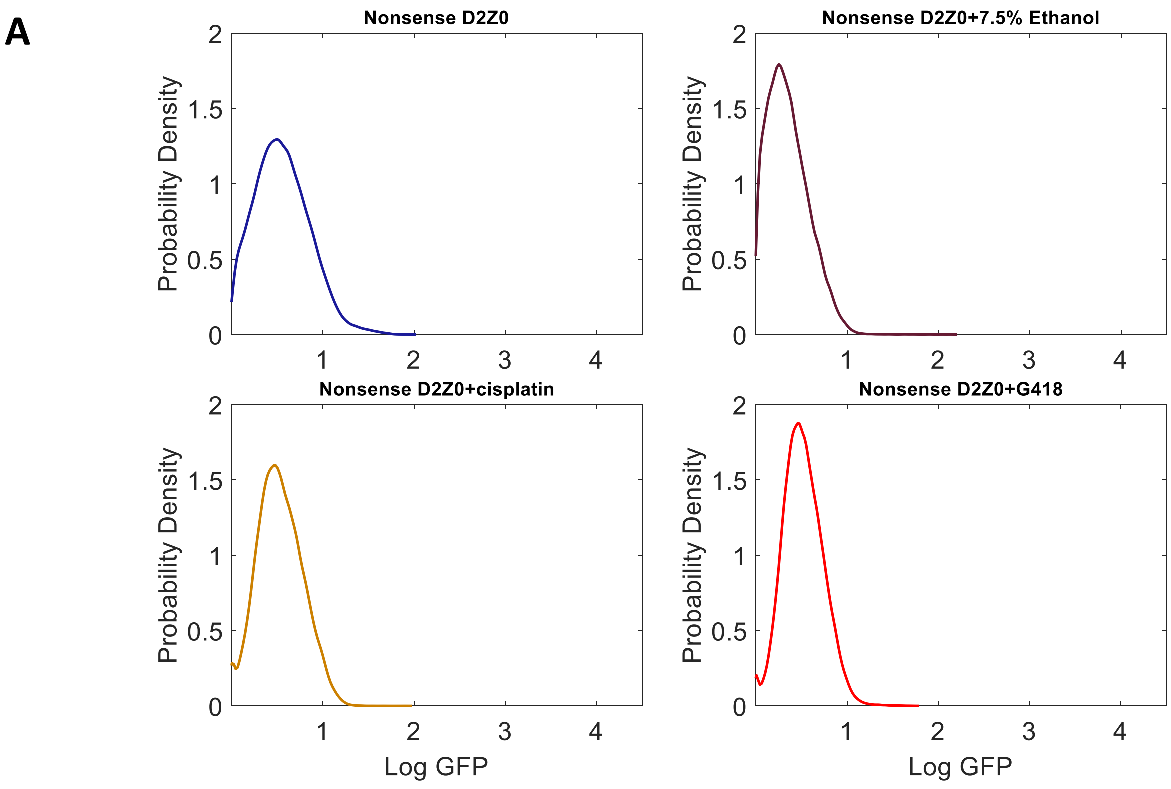

**B**

| Subpop. | # of events | Percentage | Geometric mean |
|---|---|---|---|
| **Nonsense D2Z0 g=0.278** | | | |
| Subpop. | # of events | Percentage | Geometric mean |
| High GFP | 0 | 0.00% | N/A |
| Low GFP | 109786 | 100.00% | 2.80 |
| **Nonsense D2Z0 + 7.5% EtOH g=0.071** | | | |
| Subpop. | # of events | Percentage | Geometric mean |
| High GFP | 4 | 0.00% | 54.00 |
| Low GFP | 98914 | 100.00% | 1.85 |
| **Nonsense D2Z0 + Cisplatin (40ug/mL) g=0.185** | | | |
| Subpop. | # of events | Percentage | Geometric mean |
| High GFP | 0 | 0.00% | N/A |
| High GFP | 106642 | 100.00% | 3.37 |
| **Nonsense D2Z0 + G418 (40ug/mL) g=0.184** | | | |
| Subpop. | # of events | Percentage | Geometric mean |
| High GFP | 0 | 0.00% | N/A |
| Low GFP | 102878 | 100.00% | 3.28 |

**Figure S6.** Effect of slow growth rate due to ethanol and Cisplatin on Nonsense distributions in D2Z0 at day 4. (A) Gene expression distributions. (B) Histogram statistics.

**A**

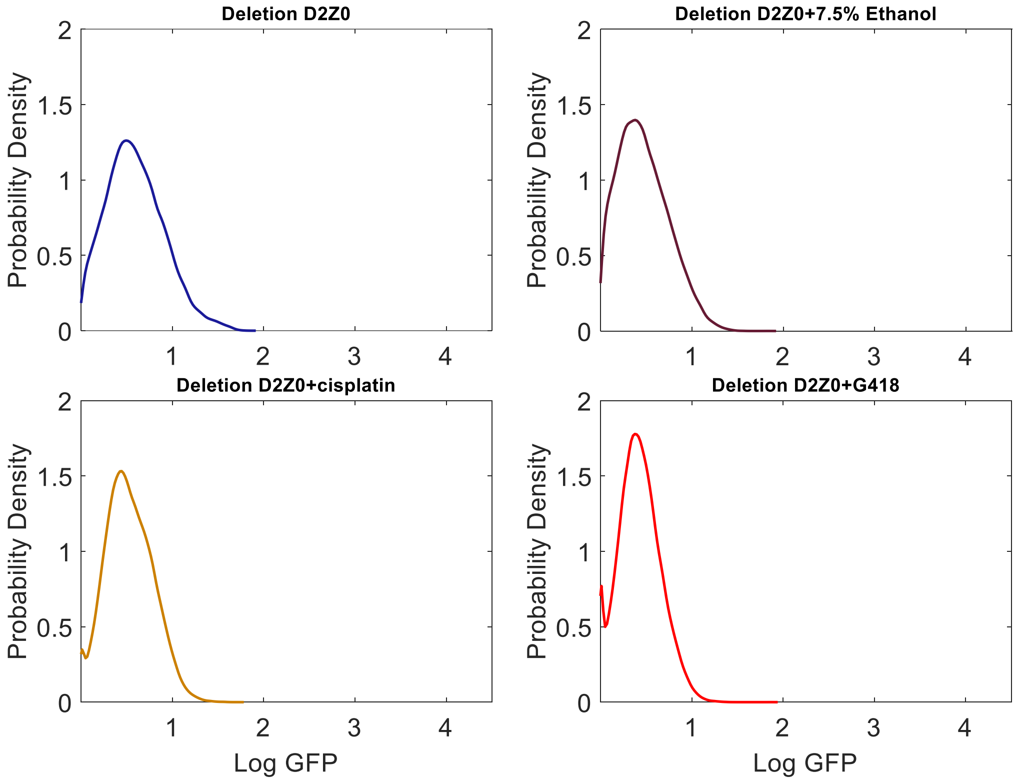

**B**

| Deletion D2Z0 g=0.287 | | | |
|---|---|---|---|
| **Subpop.** | **# of events** | **Percentage** | **Geometric mean** |
| **High GFP** | **0** | **0.00%** | **N/A** |
| **Low GFP** | **109290** | **100.00%** | **3.06** |

| Deletion D2Z0 + 7.5% EtOH g=0.130 | | | |
|---|---|---|---|
| **Subpop.** | **# of events** | **Percentage** | **Geometric mean** |
| **High GFP** | **0** | **0.00%** | **N/A** |
| **Low GFP** | **109887** | **100.00%** | **2.35** |

| Deletion D2Z0 + Cisplatin (40ug/mL) g=0.084 | | | |
|---|---|---|---|
| **Subpop.** | **# of events** | **Percentage** | **Geometric mean** |
| **High GFP** | **0** | **0.00%** | **N/A** |
| **Low GFP** | **109765** | **100.00%** | **3.35** |

| Deletion D2Z0 + G418 (40ug/mL) g=0.084 | | | |
|---|---|---|---|
| **Subpop.** | **# of events** | **Percentage** | **Geometric mean** |
| **High GFP** | **0** | **0.00%** | **N/A** |
| **Low GFP** | **109951** | **100.00%** | **2.66** |

**Figure S7.** Effect of slow growth rate due to ethanol and Cisplatin on Deletion distributions in D2Z0 at day 4. (A) Gene expression distributions. (B) Histogram statistics

A

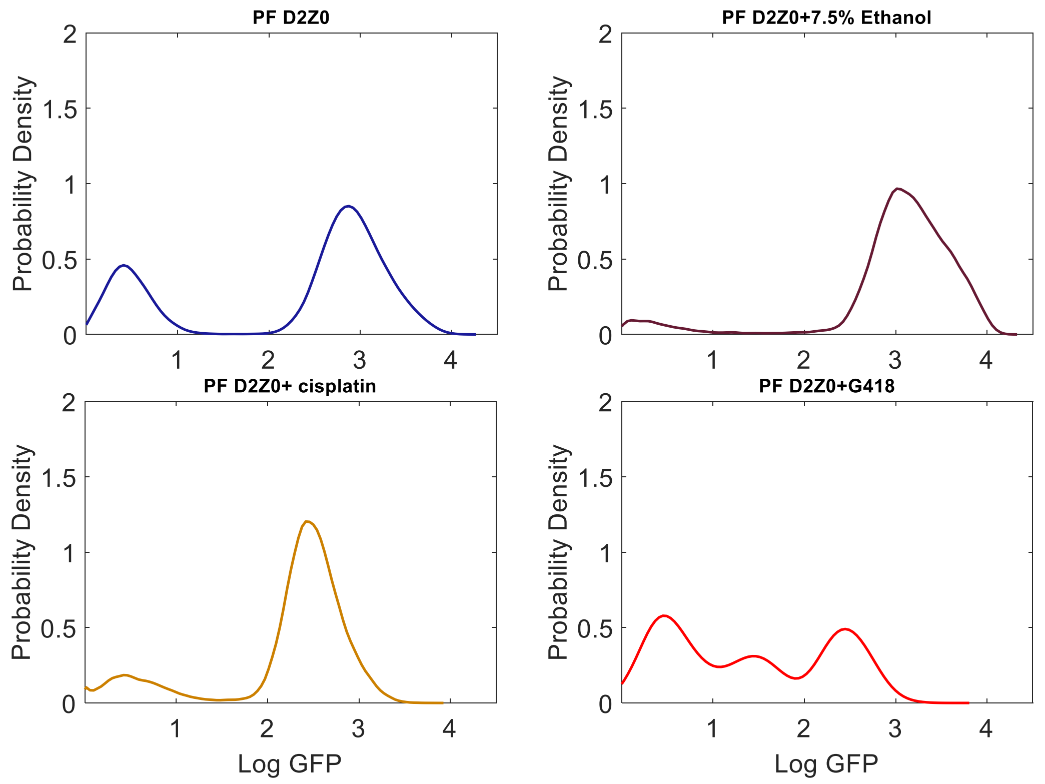

B

| PF D2Z0 g=0.146 | | | |
|---|---|---|---|
| Subpop. | # of events | Percentage | Geometric mean |
| High GFP | 77456 | 71.02% | 359.03 |
| Low GFP | 31588 | 28.96% | 2.49 |
| PF D2Z0 + 7.5% EtOH g=0.046 | | | |
| Subpop. | # of events | Percentage | Geometric mean |
| High GFP | 29216 | 90.70% | 584.10 |
| Low GFP | 2986 | 9.27% | 2.02 |
| PF D2Z0 + Cisplatin (40ug/mL) g=0.107 | | | |
| Subpop. | # of events | Percentage | Geometric mean |
| High GFP | 92993 | 84.55% | 324.14 |
| Low GFP | 16988 | 15.45% | 3.64 |
| PF D2Z0 + G418 (40ug/mL) g=0.061 | | | |
| Subpop. | # of events | Percentage | Geometric mean |
| High GFP | 38674 | 35.15% | 290.11 |
| Mid GFP | 23931 | 21.75% | 30.46 |
| Low GFP | 48024 | 43.64% | 3.34 |

**Figure S8.** Effect of slow growth rate due to ethanol and Cisplatin on ancestral PF distributions in D2Z0 at day 4. (A) Gene expression distributions. (B) Histogram statistics.

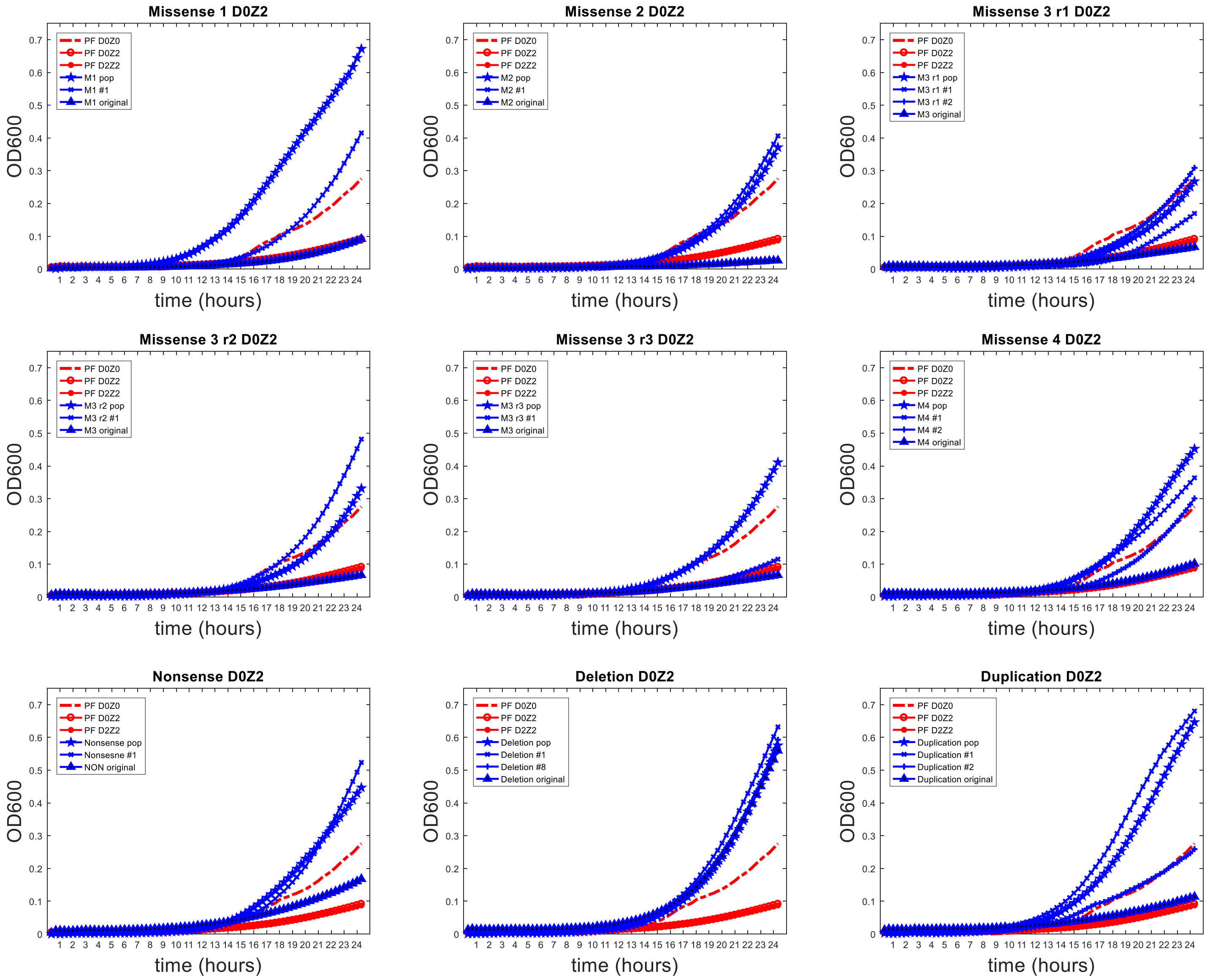

**Figure S9a**. Growth curves of original mutant populations and evolved mutant population and isolated clones. Sequenced replicate 3 of all evolved mutants is represented except for Missense 3 where replicates 1, 2 and 3 are represented.

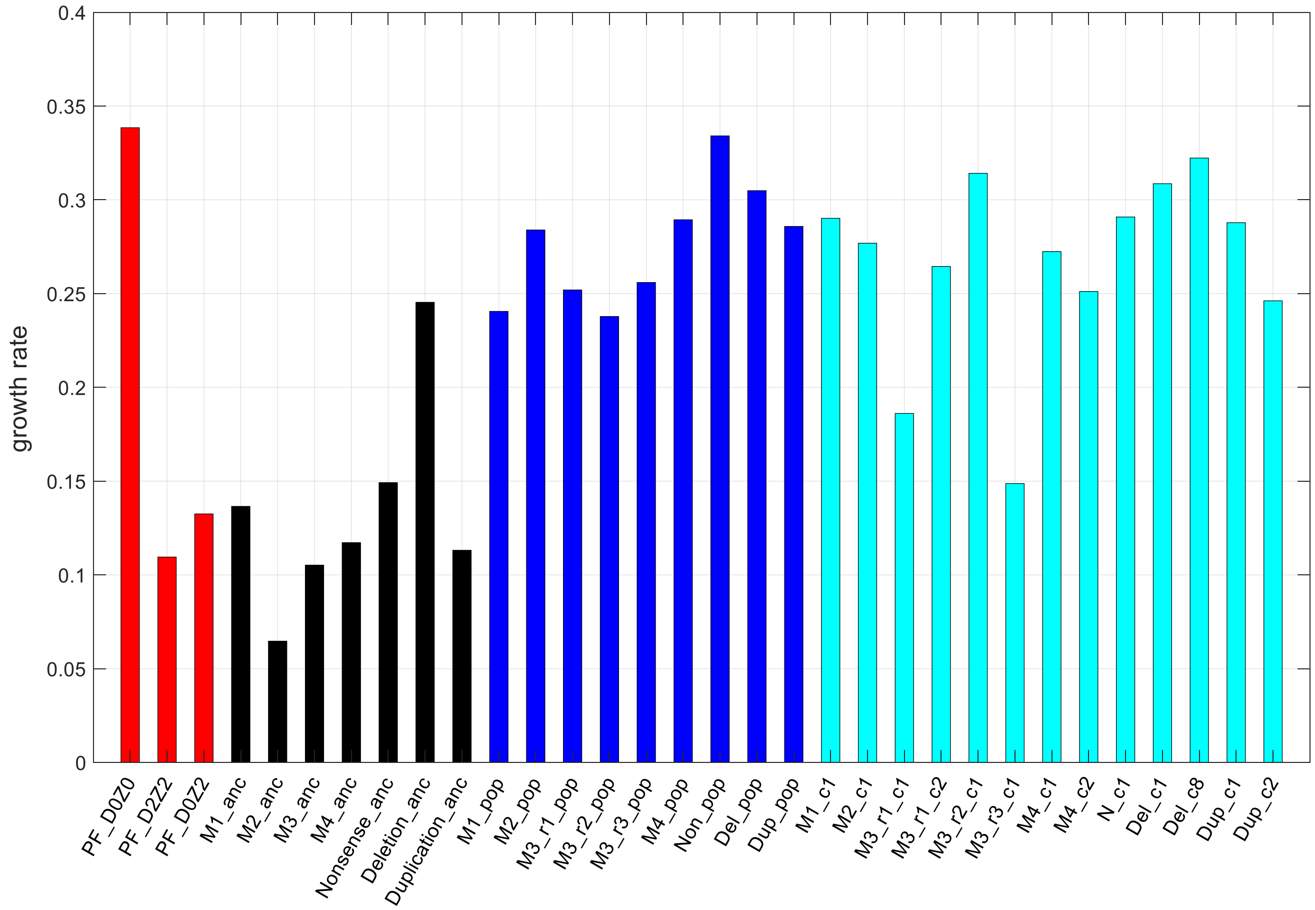

**Figure S9b**. Growth rates of original mutant populations and evolved mutant population and isolated clones. Sequenced replicate 3 of all evolved mutants is represented except for Missense 3 where replicates 1, 2 and 3 are represented.

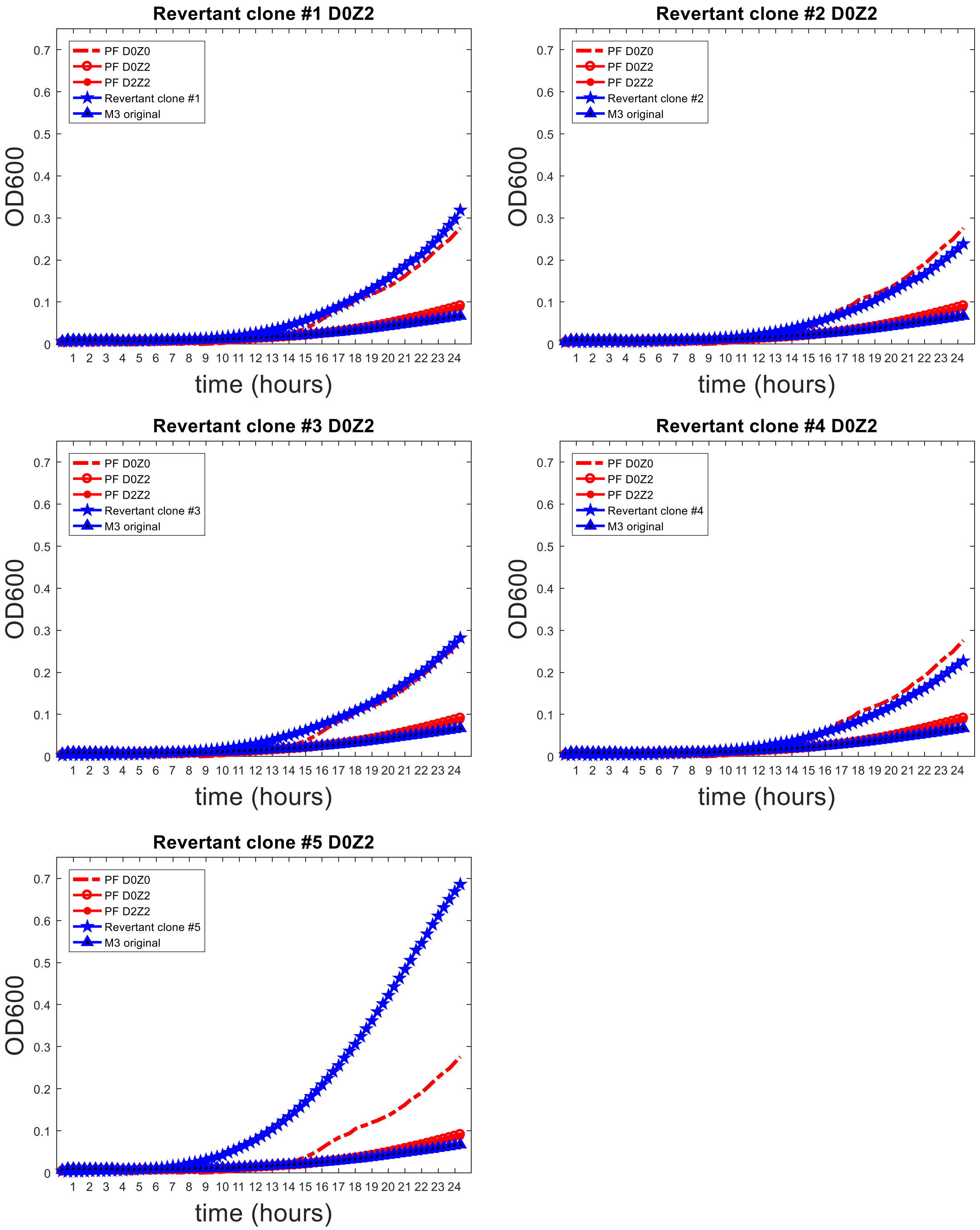

**Figure S10.** Growth curves of Revertant Missense 3 strains in D0Z2

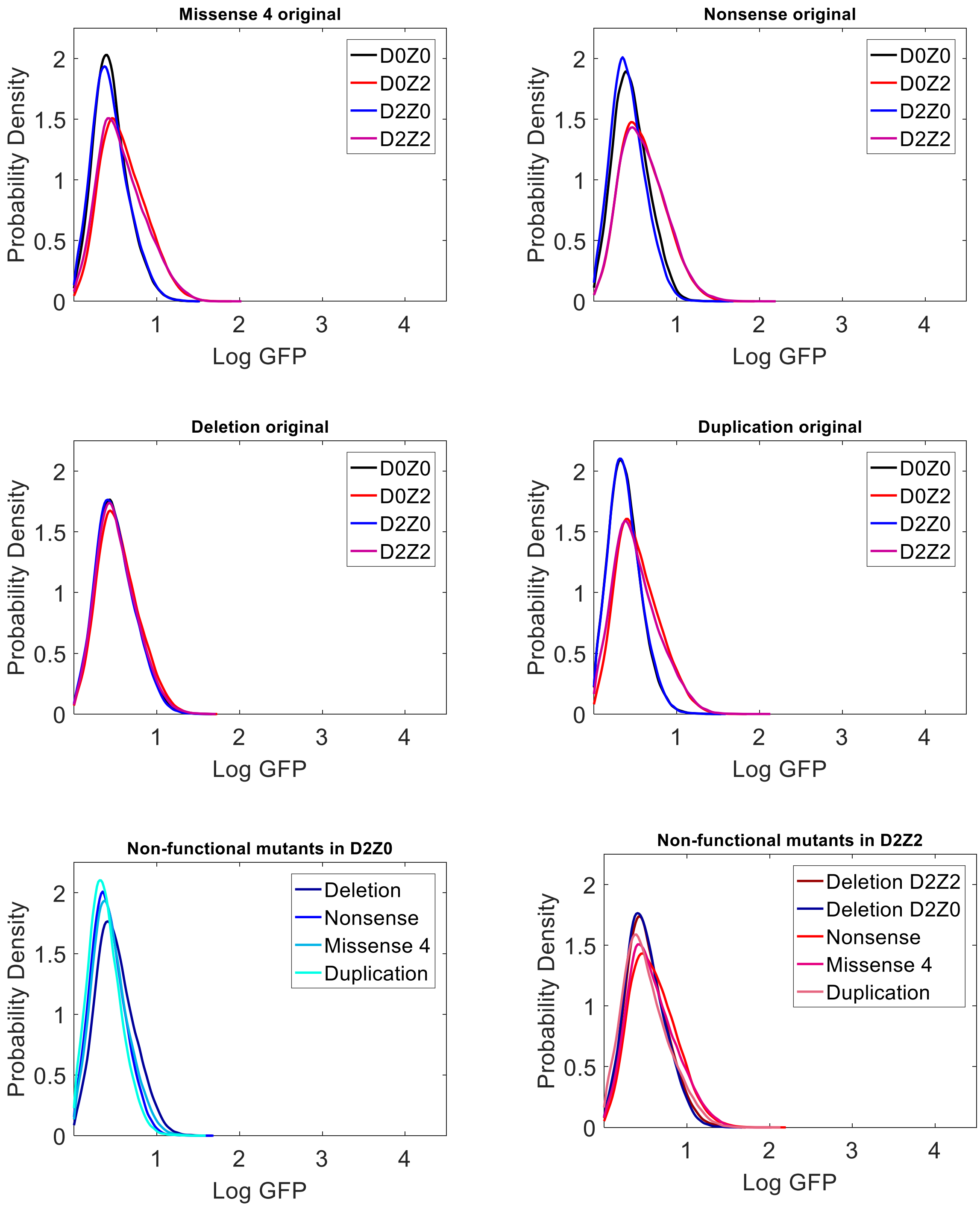

**Figure S11.** Phenotyping of original non-functional mutants. As opposed to the rest of non-functional mutants, plots show that the Deletion mutant does not undergo an upward shift in expression in the presence of Zeocin, indicating that it initially had yEGFP::ZeoR expression levels sufficient for cell survival in the presence of antibiotic.

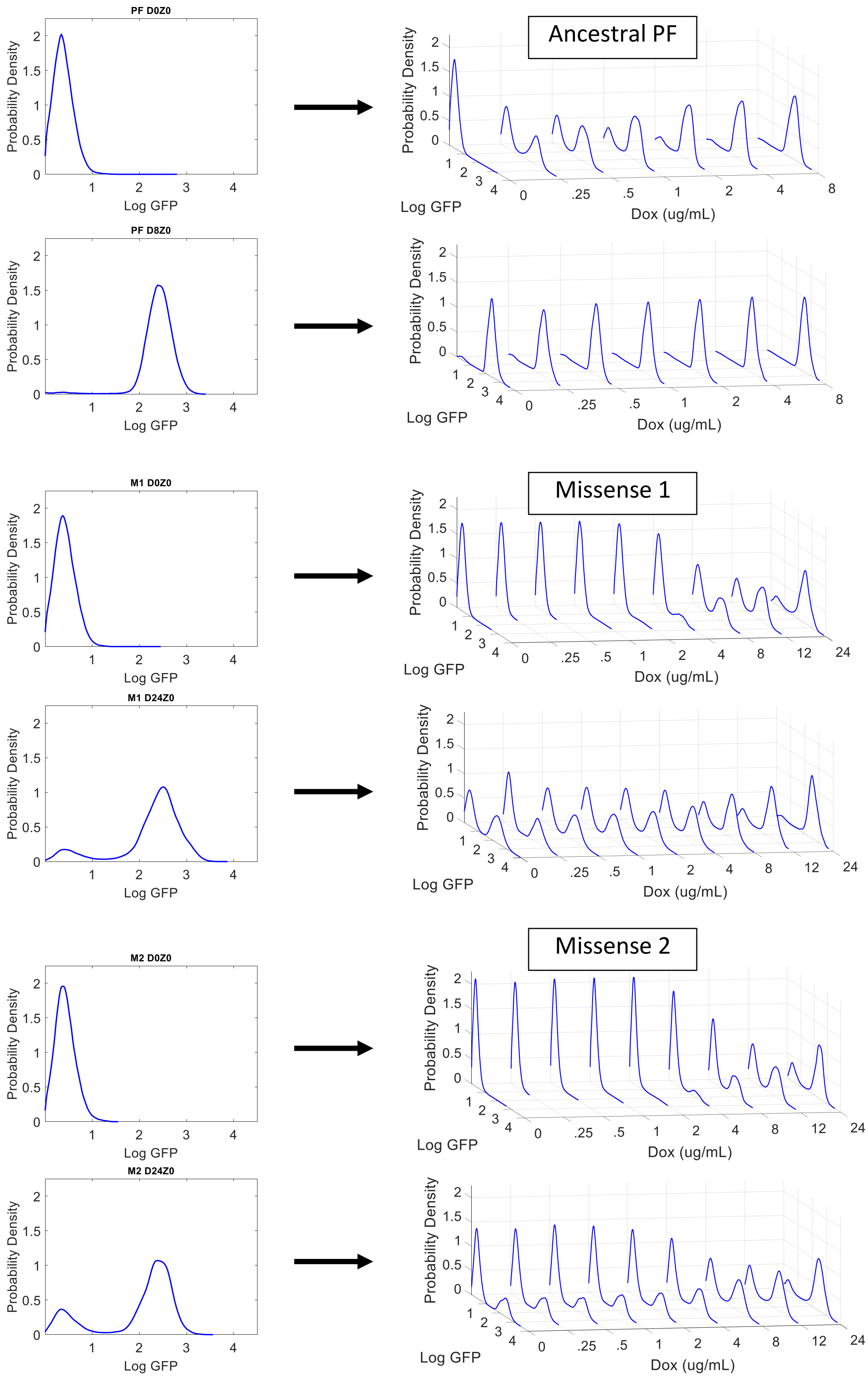

**Figure S12.** Hysteresis experiments showing the difference in Dox dose response of Missense 1 and 2 gene circuits versus the ancestral PF gene circuit within 1 day (to minimize the chance of mutations). Defining bistability based on high expressors after transfer, we obtain the ranges to be [0.05 – 8] µg/mL for ancestral PF, and [2 - 24] µg/mL for Missense 1 and 2.

### 3. Mathematical modeling

The goals of mathematical modeling are to understand: (i) wild-type PF gene circuit dynamics (e.g., why does it have two gene expression peaks above a threshold inducer concentration, why does the position of the high peak stay relatively constant, and why does the low peak not disappear even at high induction?); (ii) how the wild-type PF gene circuit broke in D2Z0, giving rise to quasi-functional, nonfunctional and dysfunctional mutants; (iii) how quasi-functional and dysfunctional mutants regained high expression in D2Z2, without any intra-circuit mutations, despite the rarity or absence of high expressors in D2Z0; and (iv) how the quasi-functional and dysfunctional mutants evolved to lose high expression without any intra-circuit mutations except for some *tetO2* operator deletions, as described in the manuscript.

#### 3.1. Deterministic (ODE) model for the wild-type PF gene circuit

We consider the following set of ordinary differential equations (ODEs) according to the drawing on the right to describe the wild-type, original PF gene circuit:

$$\frac{dw}{dt} = aF(x) - bwy - gw + l$$
$$\frac{dx}{dt} = bwy - gx$$
$$\frac{dy}{dt} = fC - bwy - (g+h)y$$
$$\frac{dz}{dt} = aF(x) - gz + l$$

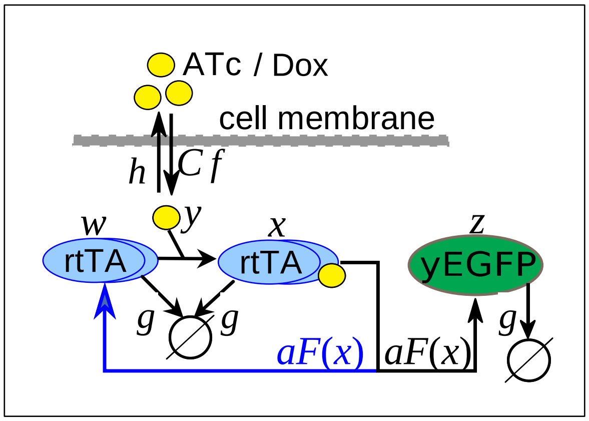

where *w* is free rtTA, *x* is active rtTA, *y* is intracellular ATc/Dox and *z* is yEGFP::ZeoR.

We assume that the promoter response function to active rtTA protein, *x* is a Hill function:

$F(x) = \frac{x^n}{\theta^n + x^n} + l$, where $\theta$ and $n$ are Hill parameters, and $l$ is a promoter leakage term.

Since total rtTA and yEGFP::ZeoR obey identical equations, their levels are identical in the model.

Next, we calculate the dose-response function, given by *z* as a function of *C* at steady state. If the system is in equilibrium, we have

$$z(C) = \frac{a}{g} F\left[x(C)\right] + \frac{l}{g}.$$

In the system of ODEs that describes the PF gene circuit there is no feedback from $z$ to the other variables. Therefore, the first three equations can be analyzed separately.

At steady state we have

$$0 = aF\left(x\right) - bwy - gw + l$$
$$0 = bwy - gx$$
$$0 = fC - bwy - (g+h)y$$

which further yields

$$0 = aF\left(x\right) - gx - \frac{g^2 x}{by} + l$$
$$w = \frac{gx}{by}$$
$$0 = fC - gx - (g+h)y$$

After substituting $y$, we have

$$0 = aF\left(x\right) - gx - \frac{g^2(g+h)}{b}\frac{x}{fC - gx} + l$$
$$w = \frac{gx}{by}$$
$$y = \frac{Cf - gx}{g+h}$$

Focusing only on the first equation:

$$0 = a\frac{x^n}{\theta^n + x^n} - gx - \frac{g^2(g+h)}{b}\frac{x}{fC - gx} + l$$

Convert this into a polynomial equation and seek real, nonnegative solutions in Matlab:

$$bg^2 x^{n+2} - g[ab + bfC + bl + g(g+h)]x^{n+1} + fbC[a+l]x^n +$$
$$+\theta^n bg^2 x^2 - g[bfC + bl + g(g+h)]\theta^n x + blfC\theta^n = 0$$

We can also pursue a graphical solution, writing the above equation as

$$aF(x)+l = a\frac{x^n}{\theta^n + x^n} + l = \frac{g}{b}\frac{b(fC-gx)+g(g+h)}{fC-gx}x \qquad \text{or}$$

$$F_L(x) = a\frac{x^n}{\theta^n + x^n} + l = \frac{g}{b}\frac{bgx^2 - [bfC + g(g+h)]x}{gx - fC} = F_R(x)$$

If $g(g+f)$ is much smaller than the other terms (notice a *singularity* at $x=Cf/g$ !!), we have:

$$F_L(x) = a\frac{x^n}{\theta^n + x^n} + l \approx gx = F_R^*(x) .$$

Essentially, the solutions are given by the intersection of a Hill-type, sigmoidal ***synthesis rate function*** $F_L$ with an elbow-shaped ***degradation rate function*** $F_R$ that is a positive-sloped line $F_R^*(x)$, except when approaching the singularity at $x=Cf/g$ from below. Any values above the singularity are non-physical.

We note that the results are robust to specific parameter values if the synthesis and loss rate curves can intersect as described below. In the next sections, we use slightly altered parameters to facilitate visualization. The actual set of parameters closely reproducing wild-type PF dynamics (including a saddle-node bifurcation at Dox=0.05 μg/mL) is the following: a=20; b=5; g=0.25; f=2; h=2.5; L=0.01; n=4; theta=0.75. The effects of the original seven mutations and then the new mutations can be modeled by parameter changes equivalent to the ones described below.

Analyzing the full equation (without making approximations, assuming $n > 2$), we obtain the following cases for the solutions (steady states) given by the intersections of red and blue curves.

**Case 1**. The line $y=gx$ misses $F_L$ from below. This occurs when $l$ is large (here, a=20; b=5; g=0.25; f=2; h=2.5; L=0.5; n=4; theta=5). The system is monostable, irrespective of Doxycycline. Only one physical solution exists, which could be low or high, given approximately by

$$x_0 \approx \begin{cases} F_L\left(\frac{Cf}{g}\right) = \frac{a(Cf)^n}{(g\theta)^n + (Cf)^n} + l; C < a+l \\ F_L\left(\frac{a+l}{g}\right) = \frac{a(a+l)^n}{(g\theta)^n + (a+l)^n} + l; C \geq a+l \end{cases} \qquad z \approx \begin{cases} F_L\left[F_L\left(\frac{Cf}{g}\right)\right]; C < a+l \\ F_L\left[F_L\left(\frac{a+l}{g}\right)\right]; C \geq a+l \end{cases}$$

The left plot below shows the dose-responses of the biochemical species *w* (free rtTA), *x* (active rtTA), *y* (intracellular inducer) and *z* (reporter) on the left. The right plot shows the right-hand side $F_R(x)$ and left-hand side $F_L(x)$ of the above equations for the system. The thick black triangle on the first (left) plot marks the Doxycycline (*C*) concentration for the plot on the right. This case illustrates a capability of the generalized system to be monostable high; this is not wild-type PF behavior. Note that total rtTA, or the sum of free rtTA *w* and bound rtTA *x* equals yEGFP, as expected due to the identical PF promoters: $z = x+w$.

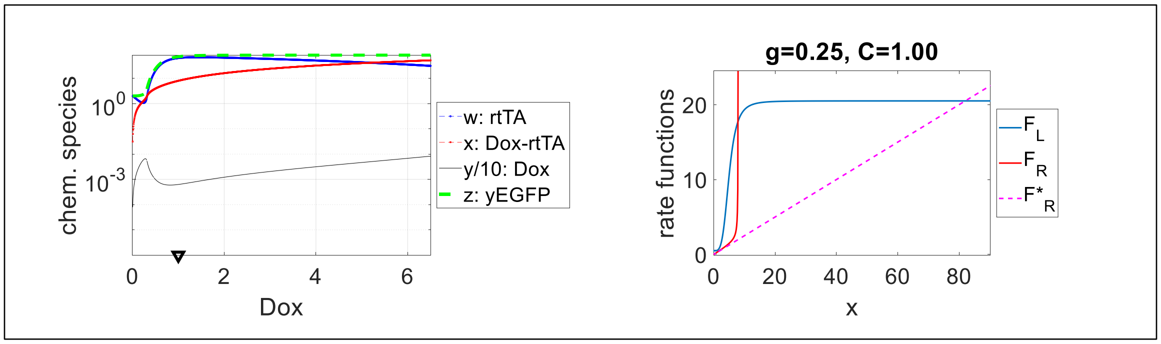

**<u>Case 2</u>**. The line $y=gx$ intersects the Hill function three times. This is the actual wild-type PF system. The number of solutions (steady states) is 1 to 3 depending on Doxycycline ($C$), as described below. The thick black triangle on the leftmost plot marks the Doxycycline ($C$) concentration for the next plots. Denoting the three intersection points of the line $y=gx$ with the Hill-type function $F_L(x)$ by $x_1$, $x_2$, and $x_3$, respectively, we have the following subcases.

*<u>Case 2A</u>*. At low Doxycycline concentrations, the system is monostable low (one intersection). This is the PF system at low induction, below the bimodality threshold, as shown below.

$$x \approx \begin{cases} \frac{Cf}{g}, \text{ if } \frac{Cf}{g} \le x_1 \\ x_1, \text{ if } x_1 < \frac{Cf}{g} \le x_2 \end{cases} \qquad z \approx \begin{cases} \frac{a}{g} F_L\left(\frac{Cf}{g}\right) + \frac{l}{g}, \text{ if } \frac{Cf}{g} \le x_1 \\ \frac{a}{g} F_L\left(x_1\right) + \frac{l}{g}, \text{ if } x_1 < \frac{Cf}{g} \le x_2 \end{cases}$$

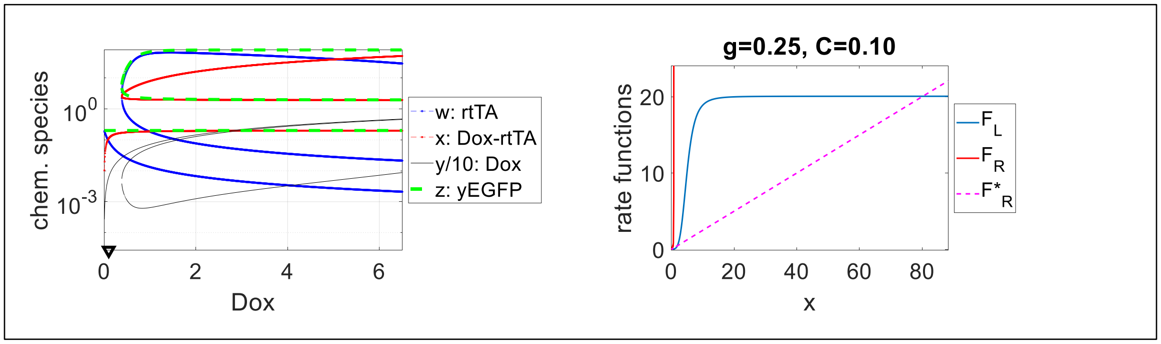

*<u>Case 2B</u>*. At medium to high Doxycycline concentrations, the system is bistable (three intersections). This is the PF system above the bimodality threshold. The system seems to remain bimodal for arbitrarily high Doxycycline concentrations.

$$x \approx \begin{cases} x_1, \text{ if } x_1 < \dfrac{Cf}{g} \le x_2 \\ x_1, x_2, F_L\left(\dfrac{Cf}{g}\right) \text{ if } x_2 < \dfrac{Cf}{g} \le x_3 \\ x_1, x_2, x_3 \approx F_L\left(\dfrac{a+l}{g}\right), \text{ if } x_3 < \dfrac{Cf}{g} \end{cases} \qquad z \approx \begin{cases} \dfrac{a}{g} F\left(x_1\right), \text{ if } x_1 < \dfrac{Cf}{g} \le x_2 \\ \dfrac{F_L\left[F_L\left(x_1\right)\right]}{g}, \dfrac{F_L\left[F_L\left(x_2\right)\right]}{g}, \dfrac{1}{g} F_L\left[F_L\left(\dfrac{Cf}{g}\right)\right] \text{ if } x_2 < \dfrac{Cf}{g} \le x_3 \\ \dfrac{F_L\left[F_L\left(x_1\right)\right]}{g}, \dfrac{F_L\left[F_L\left(x_2\right)\right]}{g}, \dfrac{1}{g} F_L\left[F_L\left(\dfrac{a+l}{g}\right)\right], \text{ if } x_3 < \dfrac{Cf}{g} \end{cases}$$

Here, to roughly capture standard PF behavior we used a=20; b=5; g=0.25; f=2; h=2.5; L=0.05; n=4; and theta=5. The behavior is shown on the plots below.

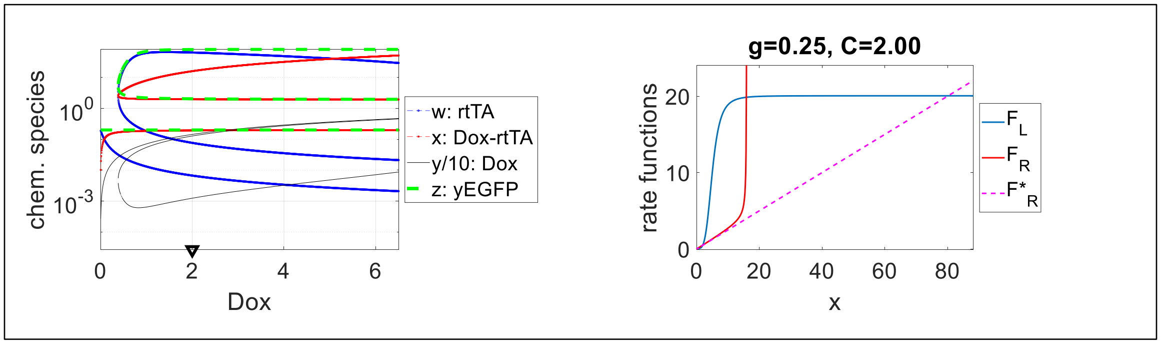

The system undergoes a saddle-node bifurcation at $Cf = gx_2$, where a new stable and an unstable node emerge. The system is bistable for any $Cf > gx_2$, thus two peaks exist even at very high Dox. The saddle-node bifurcation occurs as the function $F_R(x)$ describing the right-hand side of the equation first touches, then crosses the Hill-type function $F_L(x)$ from above.

**Case 3**. The line $y=gx$ misses the Hill function from above. This occurs when $a$ is small, and $g$ is large, for example. The system is monostable low, irrespective of Doxycycline concentration. Please note that this case just illustrates a capability of the generalized system to be monostable low; this is not standard wild-type PF behavior. Only one low-expressing physical solution exists, given approximately by

$$x_0 \approx \frac{l}{g} \Rightarrow z \approx \frac{1}{g} F_L\left(\frac{l}{g}\right) = \frac{a}{g} \frac{l^n}{(g\theta)^n + l^n} + \frac{l}{g}.$$

To obtain this behavior, we had: a=20; b=5; g=0.5; f=2; h=2.5; L=0.05; n=4; theta=25.

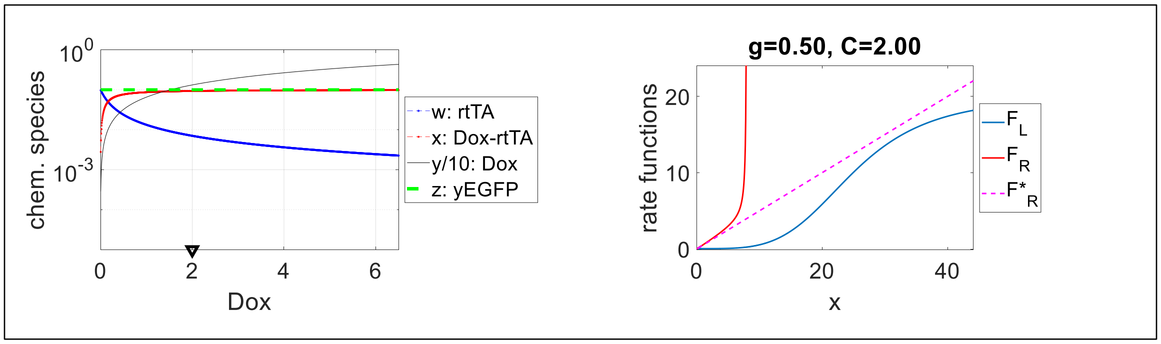

#### 3.2. Mathematical model for the nonfunctional PF mutants

Next, we ask how the PF gene circuit can be broken in nonfunctional mutants. These mutants are monostable low, regardless of Doxycycline.

One way, of course, is to completely defunctionalize rtTA, for example by flattening the Hill function (sigmoid). This means that the promoters do not respond to rtTA, as for the Deletion and Duplication mutants. These mutants cannot recover dynamically without the Hill function.

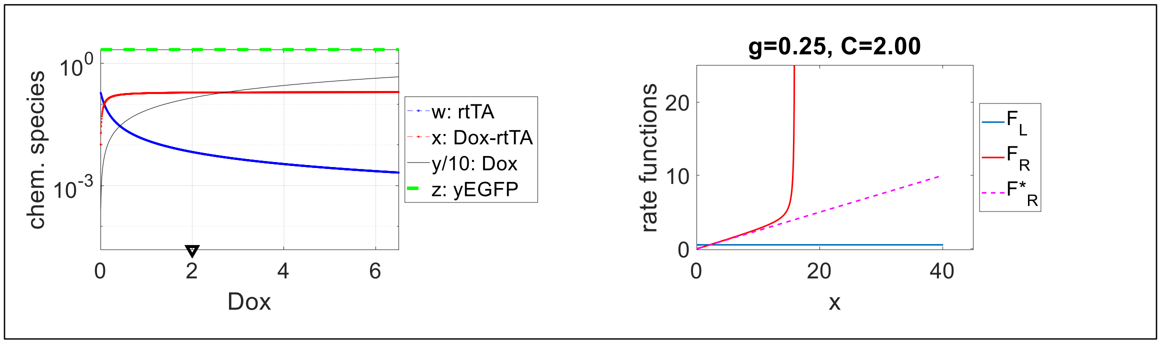

The parameters used here were: a=1; b=5; g=0.25; f=2; h=2.5; L=0.05; n=0; theta=5.

#### 3.3. Mathematical model for the quasi-functional PF mutants

Next, we seek to explain the properties and behavior of "quasi-functional" mutants that generate a high peak due to hyperinduction with Doxycycline. One way to obtain such "quasi-functional" mutants is by increasing the Hill threshold $\theta$ compared to the original PF – for example, from 5 to 25. This can certainly happen in Missense mutants if they alter the protein's binding properties to DNA. Now the magenta line $y=gx$ will still intersect the Hill function 3 times. However, due to the singularity, in D2Z0 there is only a single intersection of the Hill function with the red elbow line $F_R(x)$, hence the quasi-monostable low behavior in D2Z0. Note that the tiny experimentally observed high-expression peak can arise if the $F_R(x)$ and $F_L(x)$ curves barely intersect or if noise enables cells to access the high expression state if they are close enough.

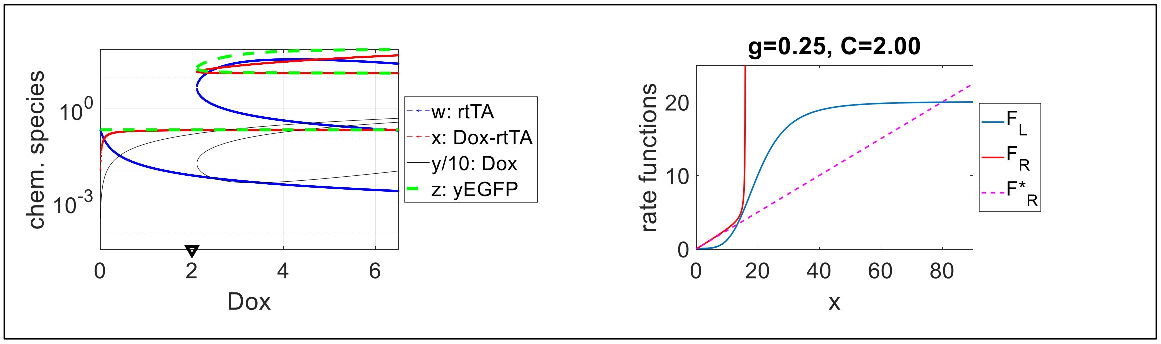

The parameter set for this mutant is: a=20; b=5; g=0.25; f=2; h=2.5; L=0.05; n=4; and θ=25.

Below we see how bistability is recovered by hyperinduction (Dox=6, D6Z0), which shifts the elbow curve's singularity rightward, allowing the red curve to intersect the blue curve 3 times.

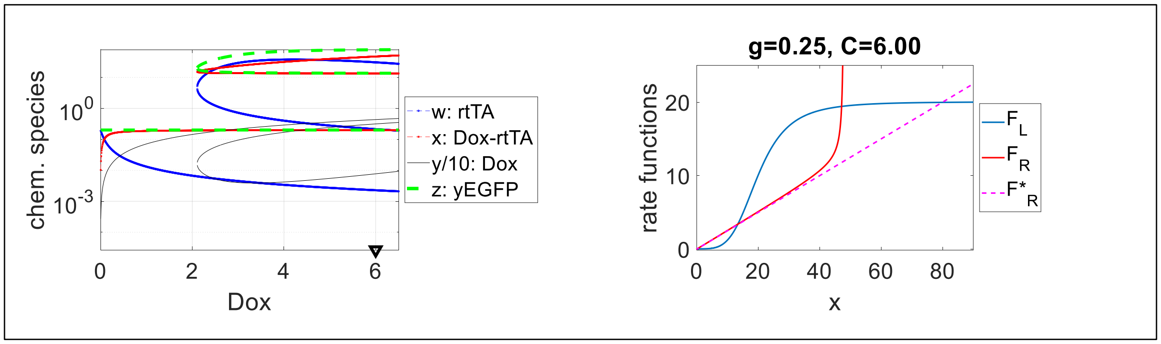

Besides hyperinduction, slow growth can also cause bistability or move the cells deeper into the bistable regime. The following plots illustrate the effect of slow growth rate (due to ethanol, Zeocin, or other factors) on the dynamics. The plots indicate that slow growth rate can convert a quasi-functional, otherwise quasi-monostable system into bistable, enabling high expression.

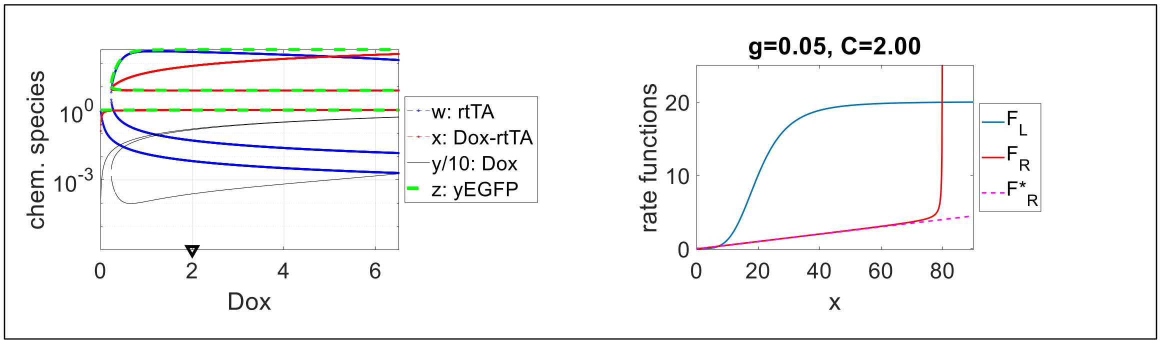

These plots imply that ethanol, Zeocin, Cisplatin or other stressors can enrich "quasi-functional" mutants in high expressors compared to pure D2Z0. So, the emergence of the high peak is due to a dynamic shift, besides phenotypic selection.

Why does the high peak diminish without any intra-circuit mutations for “quasi-functional” mutants? It is again due to a growth-related reverse dynamic shift. The bistable populations still have a low peak initially, which is hit by the drug. If any cells acquire an extra-circuit mutation that stops Zeocin from harming those cells, their growth will accelerate, returning the cells to their quasi-monostable low state, but now in D2Z2, as opposed to D2Z0. See the plots above.

#### 3.4. Mathematical model for the dysfunctional PF mutant

Finally, we seek to explain the properties and behaviors of the dysfunctional mutant Missense 3. This mutant is not hyperinducible, but still develops bistability due to slow growth. One way for this to happen is if the Hill-type sigmoidal synthesis rate function shrinks (e.g., its highest level shifts down, closer to the basal level) until the magenta and red lines miss it from above, so Missense 3 will be unimodal in D2Z0 and will not respond to any hyperinduction with Doxycycline. The parameters are: a=2; b=5; g=0.25; f=2; h=2.5; L=0.05; n=4; theta=5.

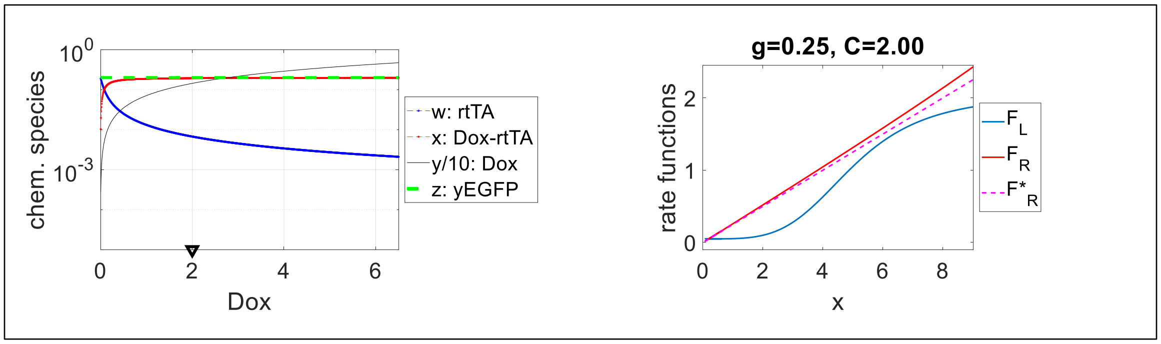

When growth decelerates due to some stressor, the red and magenta lines become less steep and can intersect the blue line 3 times, as shown below. This implies that slow growth, jointly with selection can give rise to a high expressor population, as shown below. The parameters are: a=2; b=5; g=0.05; f=2; h=2.5; L=0.5; n=4; theta=5.

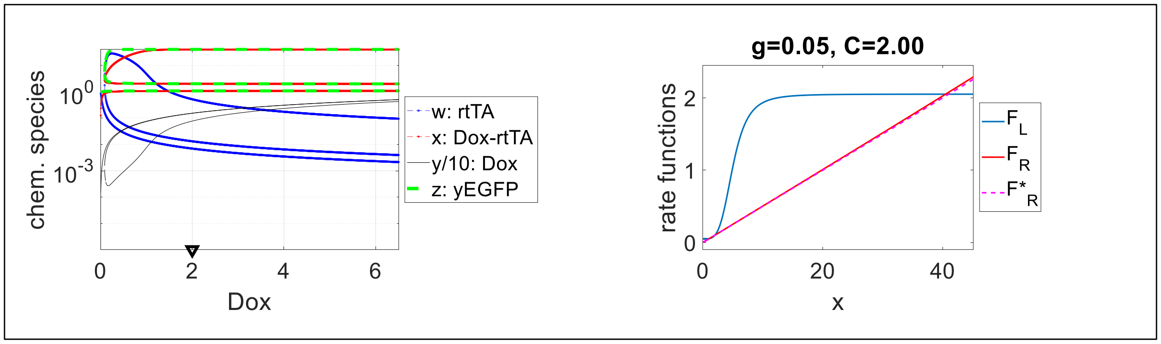

If subsequent drug-resistance mutations only speed up growth without lifting the $F_L(x)$ curve sufficiently, then the system should switch back to monostability. Indeed, we see that tendency in the population. However, some clones revert and maintain bistability. How is that possible? If extra-circuit, genomic mutations increase gene expression in general (i.e., they shift the entire $F_L(x)$ function upward) then this ensures 3 intersections that will be robust to growth acceleration. The following plots illustrate the effects of a 10-fold promoter leakage increase, from 0.05 to 0.5 while growth stays normal. The parameters are: a=2; b=5; g=0.25; f=2; h=2.5; L=0.05; n=4; theta=5.

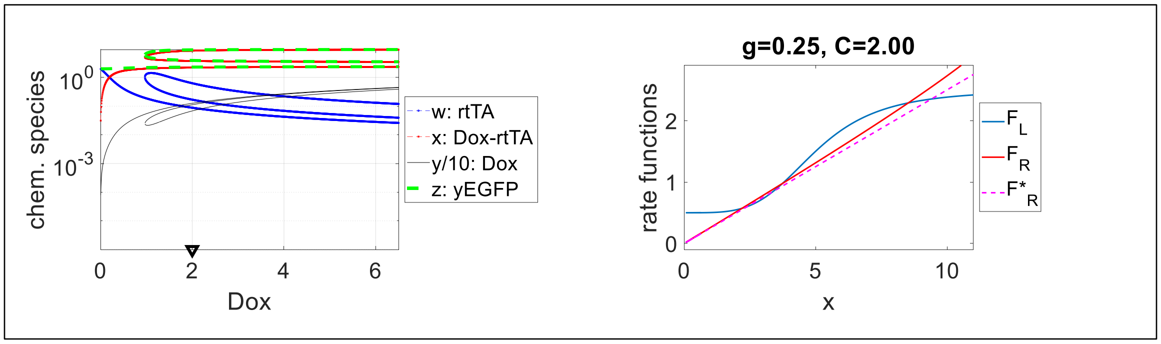

This means that the ancestral dysfunctional mutant cannot have high expression no matter how high the concentration of inducer. Yet, the dysfunctional system is not fully broken; it still has a bistable regime and it can still achieve bistability. The new mutants can maintain high expression if they can lift up the sigmoidal synthesis curve sufficiently.

For the sake of completeness, we also mention another possibility for a mutant to be dysfunctional, that is, uninducible by hyper-induction, but still inducible by slow growth. This can happen if the Hill threshold parameter $\theta$ increases by a large amount, causing line $y=gx$ to always miss the $F_L(x)$ rate function from above, regardless of Doxycycline and the singularity. We could achieve this by increasing $\theta$ from 5 to 50, which shifts the $F_L(x)$ function rightward (compare the following plots with the wild-type PF plots above). The altered parameter set is: a=20; b=5; g=0.25; f=2; h=2.5; L=0.05; n=4; and theta=50.

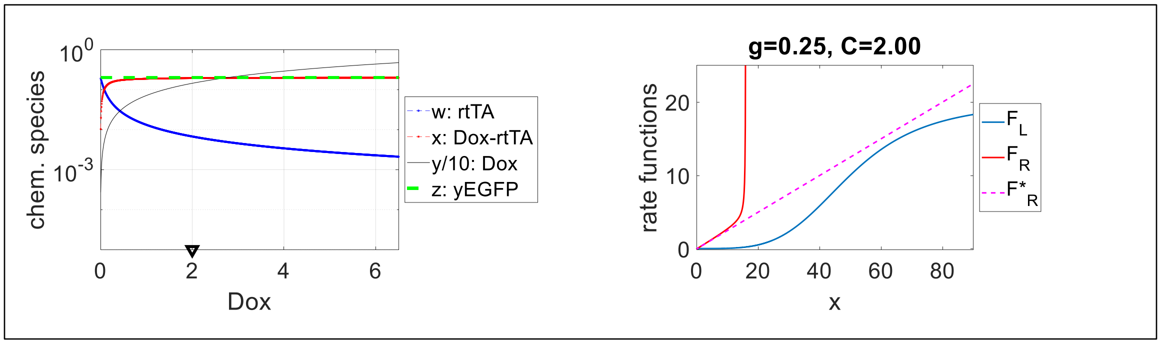

Just like the other dysfunctional mutants, such mutants could still exhibit high expression, despite being uninducible by any Doxycycline level. Slow growth, due to ethanol or Zeocin added to D2Z0 will tilt the red and magenta lines downwards, enabling them to intersect the blue $F_L(x)$ Hill-type synthesis rate function, as illustrated in the following plots. When growth speeds up again, but without the mutations affecting $F_L(x)$, bistability should be lost again. The parameter set here was a=20; b=5; g=0.05; f=2; h=2.5; L=0.05; n=4; and θ=50.

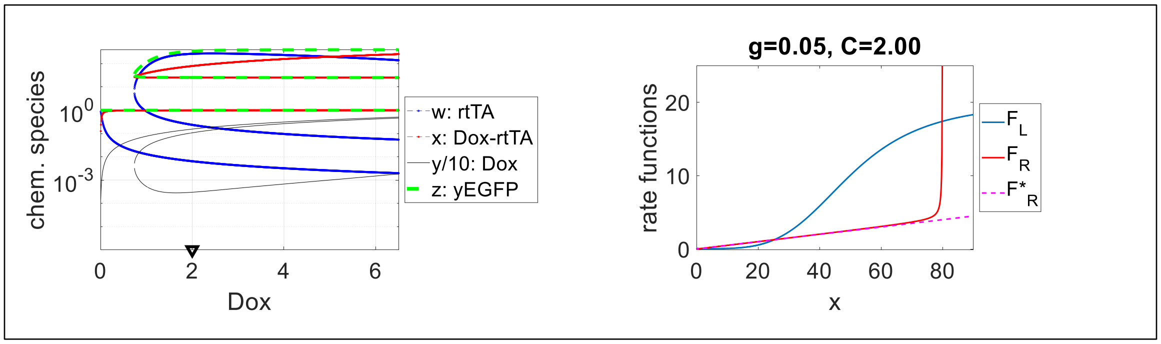

The only way to regain bistability for such mutants would be for the mutations to lower the $F_L(x)$ threshold by a large amount. This would require rtTA or PF promoter mutations. We do not think that the observed mutations are consistent with this behavior, so we consider it irrelevant to this evolution experiment.
